# De novo design of RNA pseudoknots with deep learning

**DOI:** 10.64898/2026.05.21.726960

**Authors:** Jill Townley, Wipapat Kladwang, David Baker, Hamish M. Blair, Christian A. Choe, Gina El Nesr, Andrew Favor, Eli Fisker, Daniel B. Haack, Shujun He, Jason Hingey, Po-Ssu Huang, Rui Huang, Chaitanya K. Joshi, Thomas Karagianes, Andrew Kubaney, Pietro Liò, Adamo Mancino, Jonathan Romano, Boris Rudolfs, Nicholas Spellmon, Navtej Toor, Jigyasa Verma, Vivian Wu, Zhiheng Yu, Eterna Participants, Rhiju Das

## Abstract

RNA design has been hindered by the limited accuracy of 3D structure prediction. Here, we show that intricate RNA structures can be generated with current deep learning tools through accurate de novo design of pseudoknot secondary structures. In an Eterna competition involving 57 pseudoknots, generative AI methods matched experienced human designers in solving most blind challenges, evaluated by single-nucleotide-resolution chemical mapping, compensatory mutagenesis, and cryogenic electron microscopy. AI-generated molecules with accurate secondary structures formed well-ordered 3D folds stabilized by noncanonical tertiary interactions not modeled during design. Success was guided by an RNet foundation model trained on prior chemical mapping data, suggesting that some difficult RNA design tasks may be tractable without first solving RNA 3D structure prediction.

## Main Text

Complex RNA structures underlie fundamental biological processes including translation and viral replication as well as emerging biotechnologies and medicines. Building on pioneering efforts in design by human experts (*1*), recent years have seen steady improvements in the ability to design RNA structures ranging from mRNA vaccines (*2*) to ribozymes (*3, 4*) to elaborate RNA origami (*5*), but these efforts have remained limited to design of simple RNA secondary structures, re-design of previously known RNA structures, or the assembly of a limited repertoire of well-characterized tertiary motifs. Deep learning-based design has been successful in designing RNA molecules with structures related to previously characterized RNA but with highly distinct sequences (*6–8*). However, de novo design of novel structures, while routine for proteins (*9, 10*), has not been achieved for RNA. Progress in RNA design has been limited by the poor accuracy of RNA secondary and tertiary structure prediction methods, particularly for synthetic molecules (*11–13*), for which sequences and structures of evolutionarily related molecules are unavailable.

Pseudoknots are complex secondary structures in which nucleotides in the loops of stems form base pairs with loops outside their stems (*14*). An increasing number of natural pseudoknots have been shown to function as ribozymes, riboswitches, and ribosomal frameshifting signals through the formation of complex 3D topologies with well defined stems (*15, 16*). However, novel pseudoknots have only been automatically designed for a special class of RNA origami molecules (*5, 17*) and have not been achieved with general tertiary motif assembly (*18*). In 2024, participants of the ‘OpenKnot’ challenges on Eterna (*19*) began to achieve accurate design of novel pseudoknots, as assessed and refined through experiments on millions of RNA molecules whose secondary structures were evaluated by their nucleotide-by-nucleotide reactivity to selective 2’ hydroxyl acylation read out with primer extension (SHAPE) (*20–22*). At the same time, de novo RNA design algorithms based on deep learning were proposed but remained experimentally untested in their application to novel secondary and tertiary structures (*23, 24*). These developments motivated the extension of the OpenKnot challenge to explicitly compare the design performance of these new AI algorithms to experienced Eterna participants (**Tables S1-S2**).

### Benchmarking AI design against experienced humans

Round 1 of the extended OpenKnot challenge invited six AI methods to submit designs for SHAPE experiments, in which single-stranded and conformationally flexible nucleotides are preferentially acylated at their 2’-hydroxyl groups and base paired nucleotides are typically protected and exhibit low reactivity (**Fig. 1A-E**). The 17 Round 1 secondary structures were drawn from 11 RNAs whose experimental 3D structures had been deposited in the Protein Databank (PDB) as well as six synthetic pseudoknots proposed by Eterna participants based on earlier partially successful attempts at design (**Tables S3-S4** and **fig. S1A**). For several of the 11 PDB-derived targets, the natural sequences were known to form alternative states. For example, in the Round 1 target W03, a long stem in an RNA sensing cyclic diguanosine monophosphate in *C. acetobutylicum* (cdiGMP-II riboswitch (*25*)) appeared mostly unformed in the absence of the small molecule ligand, as assessed by SHAPE (**Fig. 1D**). Therefore, even for cases with natural sequences, the challenge was to discover sequences that might stabilize the target pseudoknot for the RNA in solution without stabilizing partners. Up to 20 submissions were accepted from each AI method as well as from each Eterna participant; some methods that required input 3D backbones (**Table S2**) were not able to make designs for the 6 targets without PDB structures. Designs throughout the OpenKnot challenge were probed in the absence of small-molecule ligands so that pseudoknot formation would reflect the design rather than ligand stabilization; subsequent SHAPE measurements with ligands supported this choice (**Fig. S2, Supplementary Text**).

**Figure 1.**
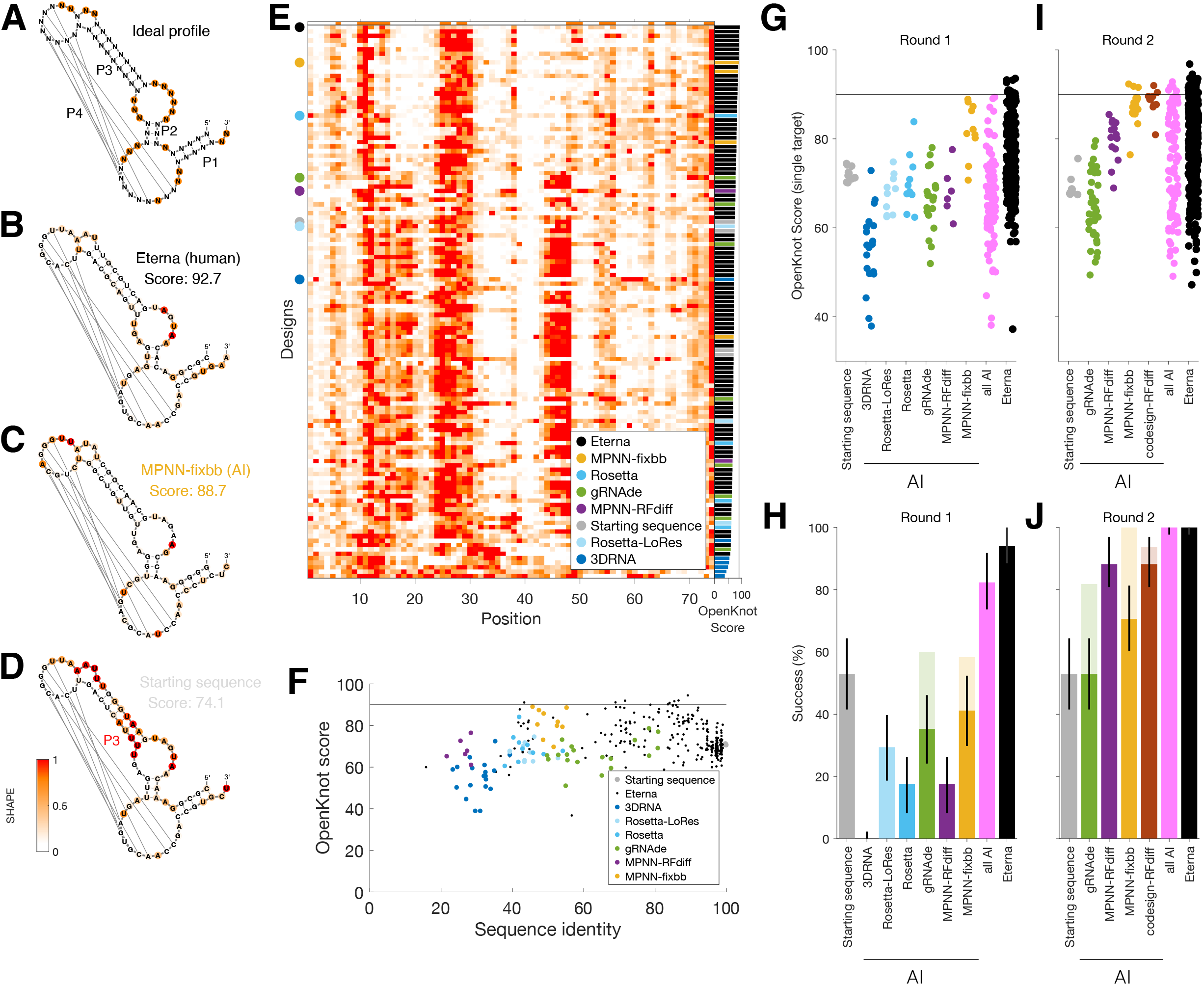
Performance on 17 starting targets. **(A-D)** Target secondary structure for W03 (‘c-di-GMP-II riboswitch’), with coloring by ideal SHAPE data (**A**) and experimental SHAPE data for top designs from (**B**) Eterna human participants, (**C**) MPNN-fixbb, and (**D**) wild type RNA (natural c-di-GMP-II riboswitch aptamer). Red label in (**D**) mark the P3 stem where SHAPE reactivity indicates flexibility of the RNA backbone, inconsistent with the target structure. (**E**) SHAPE reactivity of sequences tested for target W03 in Round 1, demonstrating a wide range of agreement with the ideal target profile (top), as parametrized by the SHAPE-based OpenKnot score (bars, far right). (**F**) Designs with low (<50%) sequence identity to the starting sequence achieved good agreement with target SHAPE profile (OpenKnot score > 90). (**G-J**) Performance summaries across design methods for **(G**,**I)** W03 OpenKnot scores and (**H**,**J**) percentage of targets W01-W17 for which success (defined here as OpenKnot scores > 90) was achieved. In (**H**,**J**), error bars reflect standard errors, and light colored bars show success rates on just targets on which designs were submitted. Performance increases were apparent from (**G**,**H**) Round 1 to (**I**,**J**) Round 2.

We observed a wide range of performance, with many Eterna designs achieving experimental SHAPE profiles with the patterns of high and low reactivity matching the target profile of unpaired and paired nucleotides (**Fig. 1B** and **1E**). The agreement of each design’s experimental SHAPE profile with its target secondary structure was quantified using the OpenKnot score, which computes the percentage of nucleotides whose SHAPE reactivity matches expectation (**Materials and Methods**); scores range from 0 to 100. A cluster of Eterna designs surpassed a cutoff of success of 90% (**Figs. 1F** and **1G**; **Fig. S3A**; **Data S1**); this cutoff was set based on the inherent experimental uncertainty in the measurements (*19*) and an error rate of ∼10% of SHAPE methods in assigning secondary structure in RNA’s with known structures (*26, 27*). Eterna participants responsible for these high-scoring designs used a combination of in-game tools, intuition, and diverse design strategies (**Supplementary Text**). Most AI methods (Rosetta, Rosetta-LoRes, 3DRNA, gRNAde, MPNN-RFdiff; **Table S2**) did not pass the score cutoff and underperformed Eterna participants in Round 1 (**Figs. 1E–H**; **Supplementary Text**). One exception was MPNN-fixbb, a message-passing neural network that designs sequences for a fixed input backbone (analogous to ProteinMPNN (*28, 29*)), which improved on average over the W03 starting sequence (**Figs. 1C, 1G**). Many of the top MPNN-fixbb designs, as well as some top scoring Eterna designs, were quite different from the starting sequence (sequence identity < 50%; see **Fig. 1F**). In addition, all designs with >90% sequence identity to the starting sequence scored under 80%, suggesting the importance of shifting away from the natural sequence.

At the completion of Round 1, a neural network RNet, trained on chemical mapping of 1M previous RNAs (*21*), was observed to give SHAPE profiles that gave simulated scores largely reproducing experimental scores, especially for poorly performing designs (**Fig. S4)**. This observation suggested that RNet modeling might allow for prospective filtering of designs. The 17 Round 1 targets were then posed again to all designers in a Round 2. Eterna designers, who were given access to RNet in the interactive interface, and AI design methods that were updated to take into account RNet (gRNAde, MPNN-RFdiff, MPNN-fixbb, and the new codesign-RFdiff; **Materials and Methods**) did notably better in this round, e.g., on the W03 cdiGMP-II riboswitch target (**Fig. 1I** and **Fig. S3B**). Over all 17 targets, all the tested AI methods and Eterna human participants were able to achieve scores above 90 on at least 80% of the targets that they designed (light colored bars, **Fig. 1J**), and there was no statistically significant difference between methods. For both Rounds 1 and 2, an alternative Z-score-based assessment modeled on procedures used in the Critical Assessment of Structure Prediction (*12, 13*) gave similar rankings (**Figs. S5A-B**).

### Generalization to novel pseudoknot targets

The results above indicated that AI designers and Eterna participants could learn from explorations in prior designs to improve from Round 1 to Round 2, on the same set of 17 targets. As a test of the generalization of the resulting methods, we carried out a Round 3 with 20 new pseudoknots as target secondary structures (**Figs. 2A-C**; **Fig. S1C**; **Table S5**). Compared to Rounds 1 and 2, fewer targets (5 vs. 11) were drawn from RNAs with experimental 3D structures in the PDB, to mitigate any effects of training set memorization for the AI methods. An additional 5 target secondary structures were drawn from Pseudobase, an expert-curated database of natural pseudoknots (*30*). The remaining 10 were curated from secondary structures proposed by Eterna participants (**Materials and Methods**). As an additional test of generality, we posed 20 additional targets with longer lengths, up to 240 nts, drawn from similarly diverse sources, in a parallel Round 4 (**Figs. 2D-F**; **Fig. S1C** and **Table S5**). To enable all methods to make designs for targets that were not associated with PDB structures, modeled 3D structures were provided for targets missing experimental structures (**Materials and Methods** and **Table S5**).

**Figure 2.**
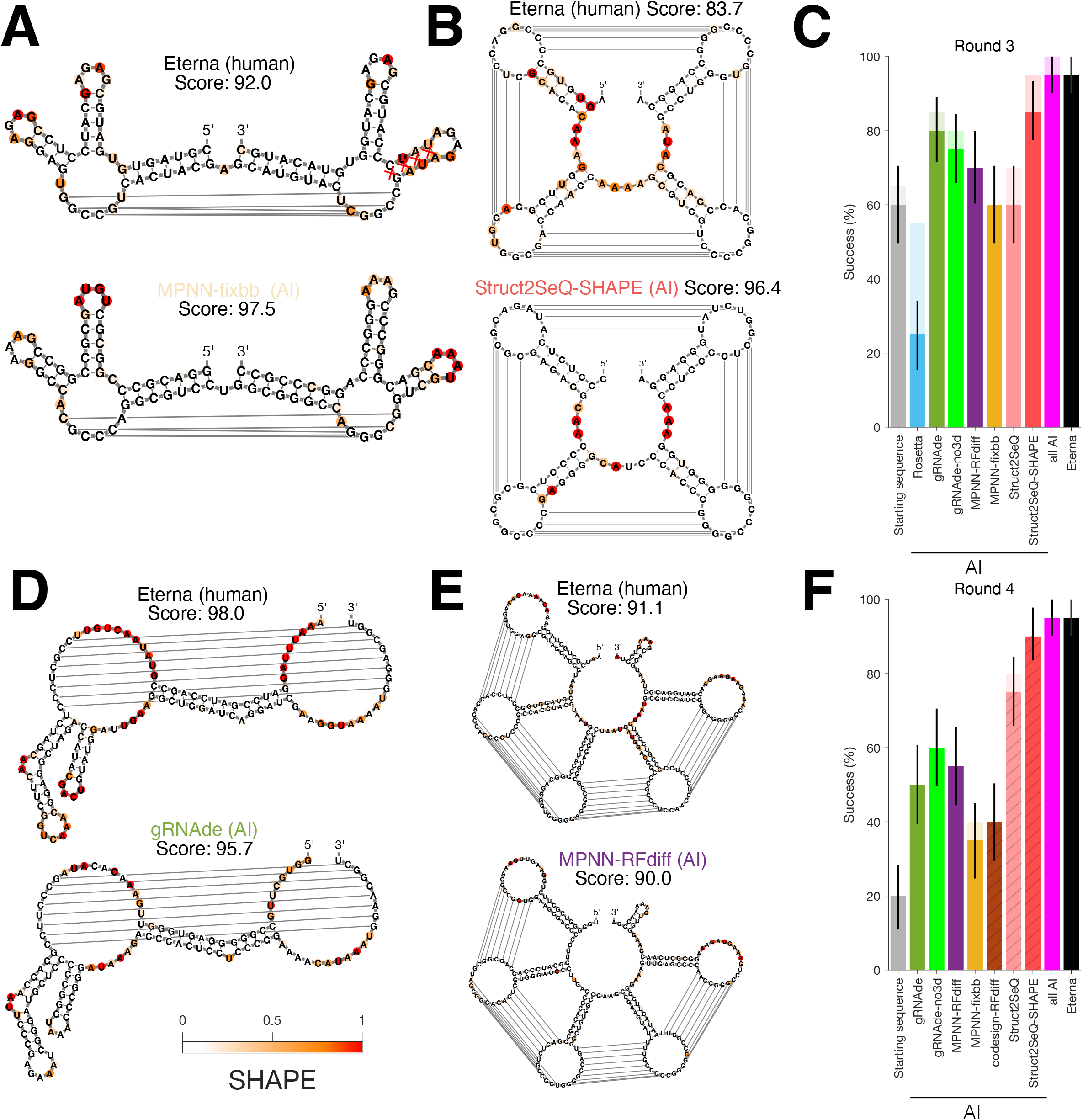
Performance on 40 new target pseudoknots. **(A-B)** Designs with highest OpenKnot score for (**A**) P20 (‘Kissing Multiloops’) and (**B**) P16 (‘AK-PK100_3’) from Eterna and AI design methods (MPNN-fixbb and Struct2SeQ-SHAPE, respectively). The Struct2SeQ-SHAPE design in (**B**) is more consistent with an alternative secondary structure (see also **Fig. S6** and **Fig. S7**). (**C**) Performance summary across 20 targets of Round 3, with lengths up to 100 nt. **(D**-**E)** Designs with highest OpenKnot score for larger Round 4 targets (**D**) Q08 (‘Rous Sarcoma Virus’) and (**E**) Q20 (‘SV_i’) from Eterna and AI design methods (gRNAde and MPNN-RFdiff, respectively). (**F**) Performance summary across 20 targets of Round 4, with lengths between 117 and 240 nt; hatched bars mark methods whose submissions were characterized in an experiment that followed release of initial Round 4 results. In (**A**,**B**,**D**,**E**), secondary structures and coloring are for the target designs and experimental SHAPE profiles, respectively. In (**C**,**F**), solid colored bars show rates of achieving OpenKnot score > 90, after designs filtered for accurate RNet-predicted secondary structures; light colored bars show rates without the secondary structure filter; and error bars reflect standard errors.

SHAPE characterization showed strong support for the performance of AI methods and Eterna participants. Each individual deep learning design method outperformed the starting sequences, whose OpenKnot scores were particularly low for the longer targets in Round 4. When grouped together, both AI methods and Eterna methods achieved OpenKnot scores of 90 in 19/20 targets in Round 3 (**Fig. 2C**) and in Round 4 (**Fig. 2F**; see also **Figs. 2C-D**). Z-score analysis also supported these conclusions (**Figs. S5C-D**). Only one target was not designable with OpenKnot score > 90 by any method or Eterna participant (P16, “AK_PK100-3”; **Fig. 2B, Supplementary Text**).

Besides the P16 target, successes included a wide variety of elaborate pseudoknots proposed by Eterna participants (e.g., Kissing Multiloops, **Fig. 2A**, and SV_i, **Fig. 2E**) as well as a pseudoknot proposed for a natural RNA with unknown 3D structure (Rous Sarcoma Virus frameshift signal; **Fig. 2D**). The success of the design methods was particularly striking for the targets with long length in Round 4, for which the starting sequences were successful in only 4 of 20 targets (the starting sequence in **Fig. 2F**). The leading AI methods included a new variant of graph neural network gRNAde that did not take 3D input as well as a method new to the challenge, called Struct2SeQ, a deep Q reinforcement learning method that was trained and guided purely by the RNet model (*31*) (**Table S2** and **Materials and Methods**). In both Rounds 3 and 4, the differences between the best method from the MPNN, gRNAde, and Struct2Seq were generally not statistically significant (*p* > 0.05; exact paired binomial test on discordant pairs, one-sided). The exception was Struct2SeQ-SHAPE which significantly outperformed the other AI methods in Round 4; this method’s designs were submitted after release of initial results, however, allowing additional development time (**Materials and Methods**).

### Compensatory mutagenesis confirms designed base pairs

The results of Rounds 3 and 4 suggested that both experienced human designers as well as AI methods had improved to the point that they could design natural and non-natural pseudoknots without progressive refinement. However, the evaluation of these designs was based on a predetermined but arbitrary cutoff of 90 for the OpenKnot score and the assumption that SHAPE measurements accurately monitor base pairing status. Caveats for SHAPE interpretation have been described (*26, 27, 32, 33*), and it is possible that nucleotides whose protection from SHAPE appeared consistent with the target secondary structure were instead forming alternative pairings. To test this possibility, we brought to bear an independent approach based on comparative mutagenesis, called mutate-map-rescue read out by high throughput sequencing (M2R-seq) (*33, 34*), illustrated for a gRNAde design of P20 (‘Kissing Multiloops’) from Round 3 in **Fig. 3A-F**. If two nucleotides form a base pair in a putative RNA structure, mutation of either nucleotide should disrupt the pair and produce SHAPE perturbations near the site of mutation (**Fig. 3A–C; Fig. S6** shows detailed data for representative designs); reactivity changes at the unmutated partner can also occur but are not always evident, since the unmutated nucleotide may remain stacked or form alternative interactions (*33, 35, 36*). However, if the two nucleotides are mutated at the same time so as to flip the original base pair (e.g., C-G to G-C), the RNA’s structure should be ‘rescued’, and the SHAPE profile should return to that of the unmutated RNA (compare **Fig. 3D** to **Fig. 3A**, and bottom and top rows of **Fig. 3E**).

**Fig. 3F** summarizes ‘rescue factors’ (*33*) for every target base pair in this P20 design. The data showed strong evidence for each stem being formed (conversely, **Fig. S7** shows a negative example, target P16). M2R-seq data were acquired across all 20 Round 3 targets for the top design from Eterna and each AI method (over 10,000 sequences, single and double mutants), and following established calibration (*32, 33*), we evaluated stem-wise recovery. Across 17 of 20 targets, at least one design achieved 100% recovery of all target stems by our M2R-seq rescue-factor criterion. For 19 of 20 targets, there was at least one AI or Eterna design with at least 80% M2R-seq stem-wise recovery (**Data S1**). With this criterion, AI methods were again competitive with Eterna (**Fig. 3G**; 90% and 75% designs successful, respectively; difference not statistically significant). The differences between AI methods were not statistically significant, except that each of the deep learning methods significantly outperformed Rosetta (*p* < 0.05; exact paired binomial test on discordant pairs, one-sided). Overall, the results from compensatory mutagenesis analysis supported the picture from the OpenKnot score analysis based on SHAPE data: like experienced human RNA designers, AI methods were able to design complex pseudoknot secondary structures, including novel targets, with consistently high accuracy.

**Figure 3.**
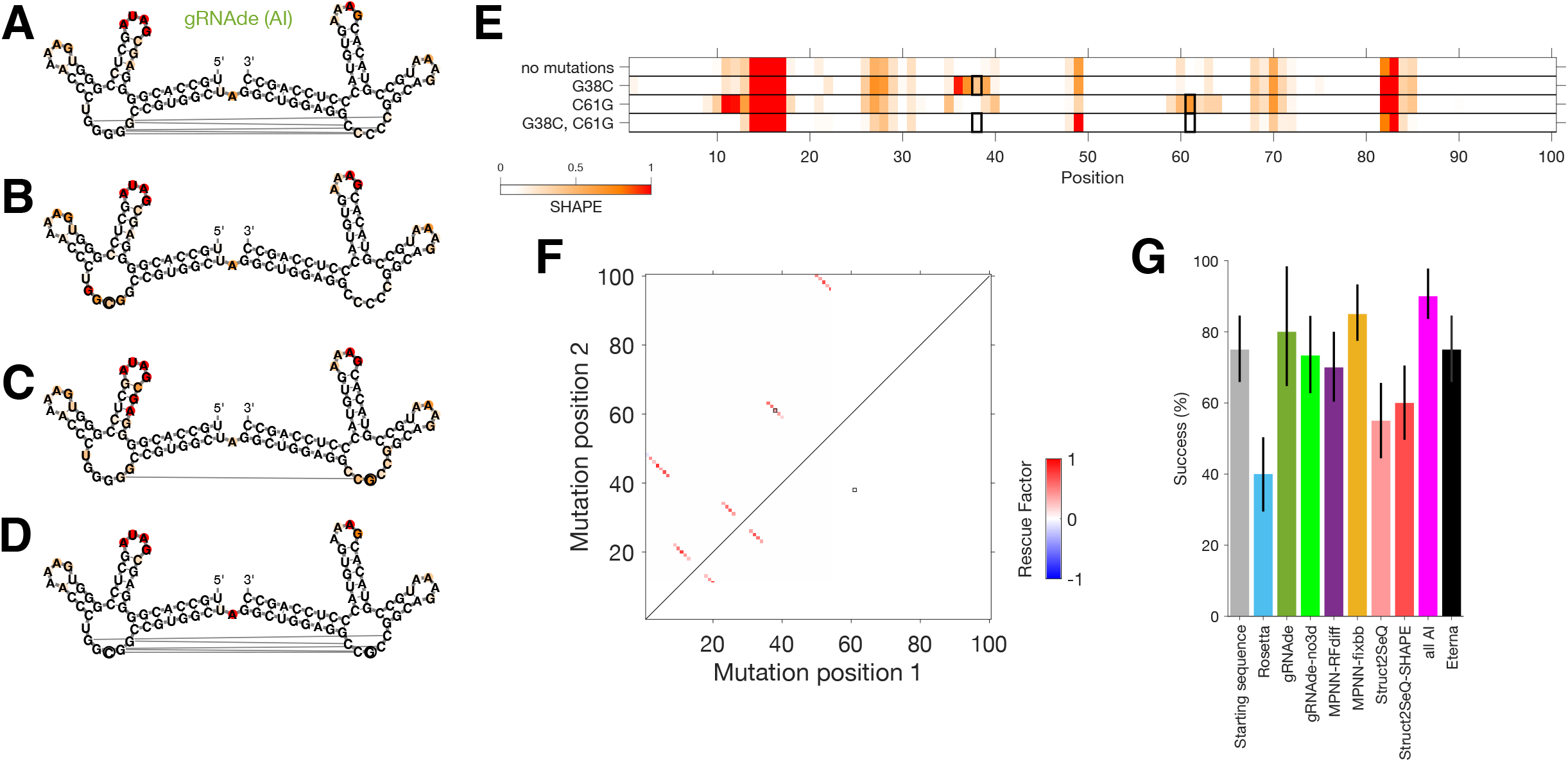
Compensatory mutagenesis (mutate-map-rescue, M2R-seq) tests accuracy of individual base pairs. (**A**) SHAPE data for gRNAde design with highest OpenKnot Score for Round 3 target P20 (Kissing Multiloops), colored onto target pseudoknot secondary structure. (**B-D**) SHAPE profiles for single mutants (**B**) G38C and (**C**) C61G showed disruptions near sites of mutations (outlined circles) that were rescued in (**D**) the compensatory double mutant which restored a 38-61 base pair. In (**A**-**D**), to aid visualization, depicted secondary structures are those modeled for the sequences by RNet-SS. (**E**) Same SHAPE profiles as in (**A**)-(**D**) stacked on each other to aid visual comparison of profile restoration upon compensatory mutagenesis. (**F**) Experimental rescue factor values for all target base pairs for the design in (**A**). Rectangles in (**E**-**F**) mark 38-61 base pair tested in (**A**-**D**). (**G**) Performance summaries across all design methods, based on the fraction of the 20 Round 3 targets in which M2R-seq verified the formation of at least 80% of target stems.

### Cryo-EM reveals novel 3D folds and noncanonical interactions

To learn if these novel pseudoknots might correspond to novel three-dimensional structures, we subjected several designs to cryogenic electron microscopy (cryo-EM). Out of the non-natural secondary structures in Round 3, we prioritized designs for the Eterna-proposed target P20 (‘Kissing Multiloops’, **Fig. 2A, Fig. 3**, and **Fig. 4A**) due to the short lengths of single-stranded linkers, which we reasoned would restrict the RNA’s 3D conformations sufficiently to enable their structural characterization via cryo-EM. Modeling in AlphaFold 3, trRosettaRNA, and other RNA 3D structure prediction algorithms also suggested that the P20 designs would form well-defined folds, though 3D modeling confidence was poor (**Fig. 4B**). To increase the visibility of the designs in cryo-EM micrographs and to break any pseudosymmetry that might preclude high resolution refinement, we embedded the top P20 designs inside a recently developed circularly permuted group II intron scaffold (*37*).

**Figure 4.**
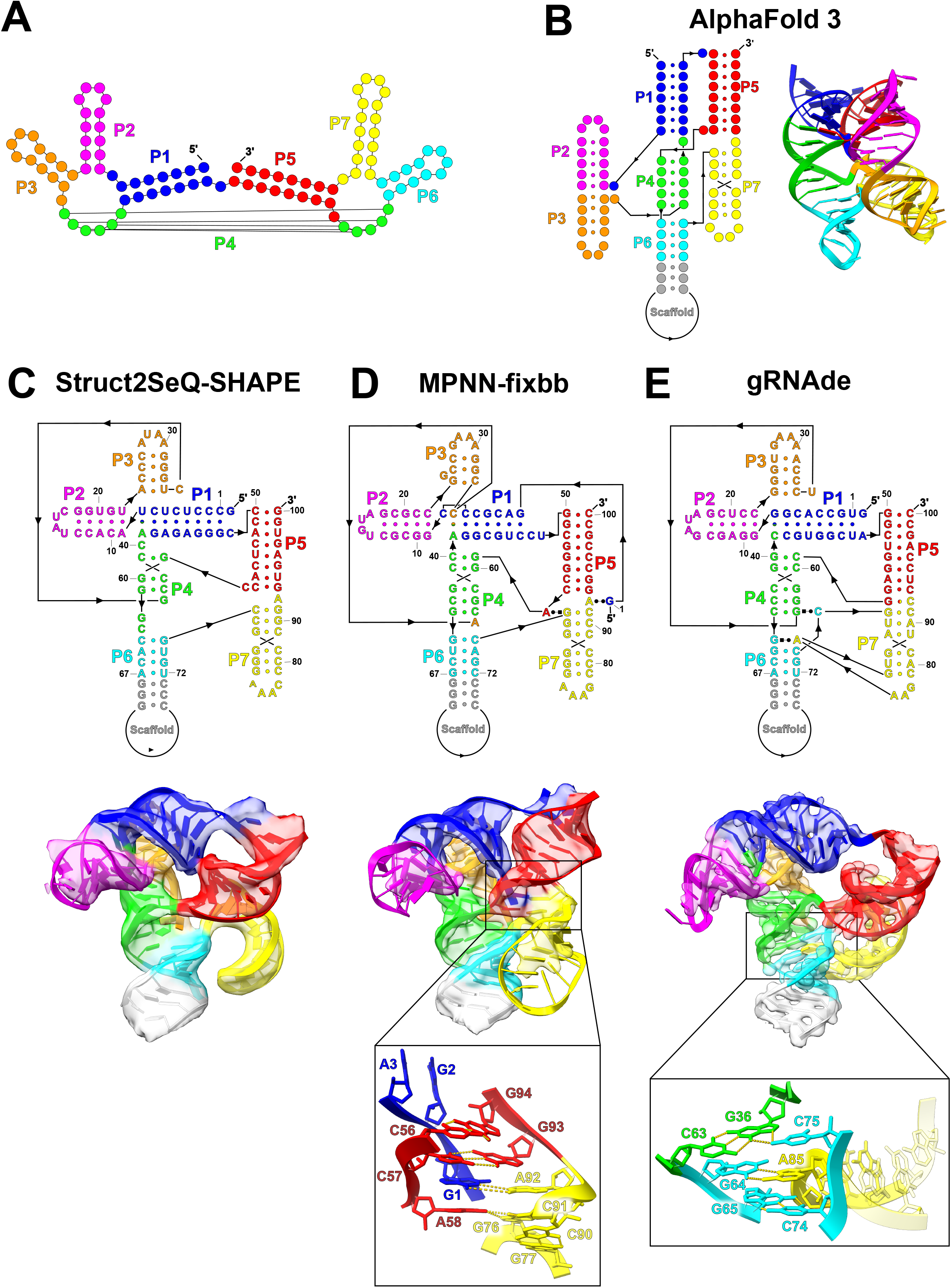
Cryo-electron microscopy of AI-designed pseudoknotted RNA. **(A)** Secondary structure of Kissing Multiloops (target P20 in Round 3), colored by stem. (**B**) AlphaFold 3 3D informed secondary structure and predicted model, colored by stem. (**C**-**E**) For the tested designs from (**C**) Struct2SeQ-SHAPE, (**D**) MPNN-fixbb, and (**E**) gRNAde, cryo-EM derived secondary structures (top), cryo-EM maps (unsharpened) and fitted coordinates, colored by stem (bottom), show high accuracy in recovering the target pseudoknot secondary structure while also highlighting distinct topologies from AlphaFold 3 prediction and noncanonical interactions (insets under (**D**) and (**E**)).

Four designs (the starting sequence, Rosetta, Eterna, and Struct2SeQ) appeared aggregated or unfolded by size exclusion chromatography during purification and were therefore not subjected to microscopy. These were also the designs that exhibited M2R-seq stem-wise recoveries less than 80%, suggesting that base pairing stability was important for retaining folded samples. The remaining four were imaged successfully (**Fig. 4C-E, Figs. S8-S11, Table S6**, and **Movies S1** and **S2**). The MPNN-RFdiff design, which gave an M2R-seq stem-wise recovery of 80%, formed a dimer under conditions used for cryo-EM (**Materials and Methods**); all seven target stems were present but P4 ‘kissing’ interactions were formed between molecules rather than within each molecule (**Figs. S8-S11, Movie S2, Supplementary Text**).

The other three designs, from Struct2SeQ-SHAPE, MPNN-fixbb, and gRNAde, had M2R-seq stem-wise recoveries of 100%. All three resolved as monomers in cryo-EM, with map resolutions of 5.2 Å, 4.8 Å, and 3.6 Å, respectively, which improved after map masking and sharpening to 4.8 Å, 4.0 Å, and 2.9 Å, respectively. In all three maps, stems P1-P7 and interconnecting linkers were clearly visible before sharpening (**Figs. 4C-E**). These maps enabled the modeling of all coordinates, initially by unbiased manual tracing that was blind to the molecular sequence. The coordinate building confirmed that all designs formed all 7 stems of the target secondary structure with high base-pair level accuracy (*F*_1_ = 0.89, 0.95, and 0.95, respectively), and that the three folds were not homologous to prior RNA structures and did not match the AlphaFold 3 models. Detailed inspection of the two highest-resolution designs revealed additional noncanonical pairs, base triplets, and tertiary interactions (**Fig. 4D-E** insets), which did not appear to be explicitly designed by the AI methods (**Supplementary Text**).

### RNA design without 3D structure prediction

Reliable de novo design of complex RNA pseudoknots is now achievable through deep learning. In the OpenKnot AI challenge, automatic AI methods became competitive with experienced human designers from Eterna in less than one year, with both kinds of approaches solving over 95% of 57 pseudoknot design targets, as tested by SHAPE mapping and compensatory rescue experiments involving approximately 50,000 sequences. Progress in the analogous problem of de novo protein design has been driven by the use of accurate computational methods for 3D protein structure prediction (*9, 10, 38*), but such tools remain unavailable for RNA (*11–13*).

Instead of relying on 3D structure prediction tools, the MPNN, gRNAde, and Struct2Seq frameworks here leveraged a model RNet trained on prior Eterna chemical mapping data that were mainly sensitive to secondary structure. Aside from this shared use of RNet, these three AI frameworks used different variants of deep learning (**Table S2**), and our results were not able to discriminate with statistical confidence which of these AI methods was better than others.

Instead, these three frameworks may be best deployed in combination, as sequence-space analysis shows that AI methods sample tight, mutually disjoint regions, providing complementary coverage of design space (**Fig. S14**). Further experimental tests of these and other emerging methods (*7, 39–41*) will be needed to clarify which frameworks will be most appropriate for these and future RNA design tasks.

The AI-generated molecules probed here by cryo-EM displayed intricate noncanonical tertiary interactions, which are critical for sophisticated functions in natural RNA molecules (*42–44*) but were not predicted *a priori*. On one hand, predictive modeling and design of such high resolution details will be important for designing novel RNA catalysts and aptamers and therefore remains an important challenge. On the other hand, the appearance of noncanonical interactions without explicit design suggest that other problems, such as re-design of machines like the ribosome and RNA polymerase ribozymes (*45, 46*) or discovery of nonredundant folds to augment RNA structure databases (*47*), might be immediately accelerated through computational design guided by accurate secondary structure prediction, without also requiring high accuracy in prospectively modeling 3D interactions in atomic detail.

## Acknowledgments

We thank C. Geary (Heidelberg) for advice on design targets; J. Nicol (Eterna) for advice on padding designs for chemical mapping; R. C. Kretsch (Stanford) for advice on target selection for cryo-EM; A. Espeleta for assisting development of 3DRNA; J. Shendure (U. Washington) and lab for sharing primer sequences for library preparation; and Nvidia DGX Cloud and NSF NAIRR Pilot (allocation NAIRR240281) for engineering support for Struct2SeQ in Round 4. Claude was used during preparation of the revised manuscript for editorial assistance with text and figure captions and with computational analyses supporting Figs. S2, S6 and S14; all AI-assisted output was reviewed and verified by the authors.

## Funding

National Institutes of Health grant R35GM122579 (RD)

National Institutes of Health grant R35GM141706 (NT)

National Institutes of Health grant R01AI165433 (SH)

National Institutes of Health grant R01GM147893 (P-SH)

National Institutes of Health grant U19AI181881 (AK)

Howard Hughes Medical Institute (RD, DB)

National Science Foundation grant 2330652 (RD)

A*STAR Singapore National Science Scholarship (CKJ)

Qualcomm Innovation Fellowship (CKJ)

University of Cambridge Dawn HPC Pioneer Project grant (CKJ)

Texas A&M X grant (SH)

Merck Research Laboratories (MRL) Scientific Engagement and Emerging Discovery Science (SEEDS) Program (PH)

## Author contributions

J.T., E.F., and R.D. designed Eterna OpenKnot targets; and J.T., J.R., and T.K. coordinated target deployment, design collection, and scoring on Eterna. C.K.J., P.L., A.F., A.K., G.E.N., S.H., D.B., P.H., C.A.C., and Eterna Participants submitted sequences. W.K., H.M.B., V.W., R.H., J.V., and R.D. designed, collected, and analyzed SHAPE experiments. D.B.H., J.H., B.R., N.T., N.S., A.M., and Z.Y. designed and carried out cryo-EM measurements and modeling. R.D. drafted the manuscript with input from all authors. All authors other than J.T.,W.K., Eterna Participants, and R.D. are listed in the byline alphabetically.

## Data, code, and materials availability

SHAPE profiles and modeled 3D structures for Rounds 1-4, M2, and M2R experiments are publicly available (*48, 49*) and at the RNA Mapping Database (https://rmdb.stanford.edu) under the following accession IDs: OK45LIB_2A3_0000 (Round 1), OK6LIB_2A3_0000 (Round 2), OK7ALIB_2A3_0000 (Round 3), and OK7BLIB_2A3_0000 and OK7BLIB_2A3_0001 (Round 4). Cryo-EM structures for designs of P20 (Kissing Multiloops) are available in the PDB under accession IDs 10ZT (gRNAde design, Mol9), 10ZU (MPNN-fixbb design, Mol14), 11EH (Struct2Seq-SHAPE, Mol13); and 11AG (MPNN-RFdiff dimer, Mol11). Cryo-EM maps are available at EMDB under accession IDs EMD-75574 (gRNAde design, Mol9), EMD-75575 (MPNN-fixbb design, Mol14), EMD-75648 (Struct2Seq-SHAPE, Mol13), and EMD-75584 (MPNN-RFdiff dimer, Mol11). Codes are available at (*50*). Materials are available upon request.

## License information. HHMI authors

### Supplementary Materials

Materials and Methods

Supplementary Text

Figs. S1 to S14

Tables S1 to S8

References (51-58)

Movies S1 to S2 Data S1

