## Supplementary Materials for "De novo design of RNA pseudoknots with deep learning"

**The PDF file includes:**

Materials and Methods
Supplementary Text
Figs. S1 to S14
Tables S1 to S8

**Other Supplementary Materials for this manuscript include the following:**

Movies S1 to S2
Data S1

### Materials and Methods

Pseudoknot targets for the OpenKnot AI challenge

The Eterna platform and its extension to pseudoknot visualization and modeling has been described previously (1, 2). Rounds 1-4 of the OpenKnot AI challenge corresponded to Eterna's Rounds 4, 6, 7a, and 7b, respectively (Table S3); prior 'free form' Eterna Rounds 1-3 challenged human participants to submit pseudoknotted RNA sequences without prescribing specific pseudoknots, as described in ref. (2). The seventeen target secondary structures for Rounds 1 and 2 were hand-selected to challenge Eterna participants, with 11 targets derived from published PDB coordinates for diverse RNA pseudoknots with lengths of 100 nt or less, one RNA origami with a modeled 3D structure (generously provided by C. Geary, Heidelberg), and five targets chosen based on the SHAPE-directed secondary structure of top scoring participants' submissions from prior Eterna 'free form' rounds. Three of these five secondary structures were synthetic and two were scanned from natural RNA. For the targets associated with PDB entries, 3D models were provided in-game to Eterna participants and as PDB-formatted files to AI design groups; for targets without PDB entries, designs for some AI methods were not submitted (gRNAde and MPNN-fixbb; see below). Within the Eterna interface, the Round 1 puzzles provided interactive feedback with the ThreshKnot algorithm run with base pair probabilities predicted by EternaFold (*51*, *52*). In subsequent rounds, the Eterna interface provided interactive folding through the RNet-SS neural network model as well as an expected $F_1$ score predicted from this model (*21*). Experimental OpenKnot scores for Rounds 1 and 2 were provided to participants after each round and before the subsequent round.

Targets for Rounds 3 and 4 had no overlap with Rounds 1 or 2. For Round 3, five structures each were chosen from PDB (3) and Pseudobase (PKB) (4) sequences characterized in a prior 'Pseudoknot detective' round of Eterna; to help ensure designability, sequences were chosen whose SHAPE profiles better matched pseudoknot predictions compared to non-pseudoknot predictions generated by a diverse set of secondary structure modeling packages (ensemble OpenKnot Score above 80; see (2)). For the synthetic structures in Round 3, five were selected from earlier SHAPE results as likely forming target pseudoknots, and five were selected by Eterna participants during a special sprint design challenge seeking unusual pseudoknots whose designability was not known. For Round 4, targets with lengths longer than 100 nt, the maximum length in Rounds 1-3, were collected. These Round 4 targets, with lengths from 117 to 240 nt, involved six PDB entries, two PKB entries, one RFAM (5) entry, and a natural sequence submitted by participants chosen from an early free-form Eterna competition. Five synthetic submissions that scored >68 were also chosen along with five untested sequences from the sprint design challenge. The pseudoknotted secondary structures for these targets are provided in Tables S4 and S5 and visualized in Fig. S1. For Rounds 3 and 4, to encourage submissions from all design methods, a 3D model of each structure was provided in-game to Eterna participants and as PDB-formatted files to AI designers. Where a complete PDB model did not exist, a predicted 3D structure was generated by the trRosettaRNA server (6). In cases where the trRosettaRNA prediction contained unrealistic bonds, a 3D model was generated by the FARFAR2 server (7). In addition, alternative 3D structures for each target were generated in RFDpoly and made available to the gRNAde developers to test the utility of this broader set of inputs.

Chemical mapping experiments

*Synthesis of RNA libraries*

Designs were collected from Eterna participants as well as AI design groups through the collection interface in the Eterna game (links at Table S3). Designs were padded to uniform lengths (100 for Rounds 1-3; 240 for Round 4), appended with hairpins with unique sequences as barcodes, and then prepended and appended with constant primers (listed in Table S7) as in earlier OpenKnot rounds (2) , using the Fast Library Design package, available at https://github.com/hmblair/fld. As controls, 10 replicates of the starting sequence for each target were included in each library with 10 distinct barcode hairpins. Synthesis of libraries was carried out by Twist Bioscience (South San Francisco, CA) and high throughput SHAPE chemical mapping was carried out as in ref. (2), with adaptations to enable cost-efficient sequencing with Ultima Genomics (Fremont, CA), as follows.

Libraries for Rounds 1-4 (referred to internally as OpenKnot 4, 6, 7a, and 7b) were synthesized by Twist Bioscience (South San Francisco, CA). Libraries were amplified through emulsion PCR, based on mixing of oil phase and aqueous phase. Reactions included 300 μL of oil phase, mixed as follows: 80 μL of ABIL EM90 (Evonik), 1.0 μL of Triton X-100 (Sigma Aldrich T8787), 1919 μL of mineral oil (Sigma Aldrich M5904). The oil phase mixture was chilled on ice for 30 minutes and transferred to a cold glass vial. A magnetic stir bar (Sigma-Aldrich, cat. no. Z329061) was placed into the glass vial and the stir rate set to 1000 rpm for 5 min before adding aqueous phase. The aqueous phase for the emulsion PCR was prepared in a 75 μL volume, containing the Twist library template 4 μL at concentration of 1 ng/μL, 3 μL of 100 μM "Eterna" forward primer, 3 μL of 100 μM Tail2-RC primer, 1.875 μL of bovine serum albumin (20 mg/mL, Thermo Fisher B14), 37.5 μL of 2X Phire Hot Start II PCR Master Mix (Thermo Fisher F125L), and RNase free $H_2O$ was added to final volume of 75 μL. The oil phase was stirred at 1000 rpm for 5 min, then 10 μL droplets of aqueous phase were added into oil phase with a pipette 5 times with 10 seconds waiting period between each addition. The emulsion mixture was stirred for 10 minutes and transferred into PCR tubes for thermocycling. The thermocycler setting was 98 °C for 30 seconds; then 98 °C for 10 seconds, 55 °C for 15 seconds, and 72 °C for 40 seconds, repeated for a total of 42 cycles; 72 °C for 5 minutes, and 4 °C hold. The emulsion PCR product was purified using QIAquick PCR Purification Kit (QIAGEN, #28104).

The RNA library was synthesized by in vitro transcription (TranscriptAid T7 High Yield Transcription Kit, Thermo Scientific #K0441) using the emulsion PCR product as the template. Each in vitro transcription reaction mixture contained 8 μL 25 mM nTPs, 4 μL of 5X TranscriptAid buffer, 2 μL of TranscriptAid Enzyme Mix, and 6 μL of 50 - 120 ng/μL emulsion PCR DNA. The reaction was incubated at 37 oC for 3 hours, and another 15 minutes after 2 μL of DNase I was added. The RNA was purified using RNA Clean & Concentrator-25 columns (Zymo, cat. no. R1017). RNA was eluted with RNase free $H_2O$ in 45 μL.

*SHAPE modification*

For Rounds 1-3, to chemically modify the RNA, the RNA was first denatured at 90 oC for 3 minutes after mixing 12 μL of 1 M Na-Bicine pH 8.5, 12 μL of 1 M KOAc, 18.6 μL of water, and 9.4 μL of 34 μM RNA. After the RNA cooled down to room temperature, 8 μL of 100 mM $MgCl_2$ was added to the mixture to refold the RNA at 50 oC for 30 minutes. The RNA library was modified with 20 μL of 2A3 ( (2-Aminopyridin-3-yl) (1H-imidazol-1-yl) methanone, TOCRIS #7376), freshly resuspended in anhydrous DMSO to concentration of 400 mM, giving reactions with final volume of 80 μL. No-modification control reactions were also prepared with DMSO added instead of 2A3. RNA libraries were modified at room temperature for 25 min. The reactions were quenched with an equal volume (80 μL) of 1M DTT. The samples were purified using ethanol precipitation, by adding 1/10 volume of sodium acetate (pH 5.5) to the sample, mixing well, adding 3.3 volumes of cold 100% ethanol and mixing again thoroughly. The sample was incubated on dry ice or at −20 °C for 30 minutes, centrifuged at 21,000 × g for 30 minutes at 4 °C in a microcentrifuge; then, supernatant was removed, and the precipitated pellet was washed with 1 mL of cold 70% ethanol and centrifuged at 21,000 × g for 5 minutes at 4 °C; and the supernatant was again removed and re-washed. The pellet was dried by leaving the microcentrifuge tube open for 5 minutes and resuspended in 47 μL of RNase free $H_2O$. For Round 4, the 2A3 stock concentration was reduced from 400 mM to 133 mM to ensure lower per-nucleotide modification rates on longer RNAs, and all reaction volumes were scaled down by 4 to help preserve the sample.

For Rounds 1-3, in reaction volumes of 200 μL, 45.0 μL of the 2A3 modified RNA (or unmodified controls), was mixed with 20.0 μL of dNTPs and 15 μL of 12 μM of Tail2-RC primers, and 89.0 μL of RNase free $H_2O$, incubated at 65 °C for 10 min and cooled on ice for 1 min, before adding 17.8 μL SSII reverse transcriptase enzyme premix (Thermo Fisher #18064022). The enzyme premix contained 40 μL of 5X First Strand buffer, 10 μL of 100 mM DTT, 12 μL of 100 mM $MnCl_2$, and 27 μL SSII reverse transcriptase enzyme (200 U/μL). The reverse transcription reaction was incubated at 42 °C for 3 hours, and the enzyme was inactivated by incubation at 95 °C for 1 min and addition of 65 μL of 100 mM EDTA. To degrade RNA, 200 μL of 0.4 M NaOH was added to each reaction and followed with an incubation at 90 oC for 3 minutes and cooled on ice for 3 min. To neutralize the reactions, 120 μL of acid quench mixture (7 mL stock made by mixing 2 mL of 5 M NaCl with 2 mL of 2 M HCl and 3 mL 3 M NaOAc) was added.

The cDNA was purified by ethanol precipitation as above and resuspended with 42 μL of RNase free $H_2O$. This cDNA was purified on gels with 10% Polyacrylamide in 7 M Urea and 1X TBE, using Criterion cassette (Biorad) to run electrophoresis at 18 W for 45 min, staining with SYBR Gold, and visualization with blue light with Dual LED Blue/White Light Transilluminator (Thermofisher part no. LB0100). Excised gel bands were purified through ZR-small-RNA PAGE recovery (Zymo Research, cat no. R1070), with final elution in 40 μL of RNase free $H_2O$.

For Round 4 libraries, the reverse transcription reactions were carried out as above, but scaled down to 40 μL volumes instead of 200 μL volumes; and cDNA was purified with RNA clean & concentrator kit (Zymo, cat. no. R1013) and elution in 15 μL of RNase free $H_2O$ and then gel cut using 2% E-gel Agarose EX gels and Qiagen MinElute PCR purification kits, with purified cDNA eluted in 15 μL of Qiagen EB buffer.

6.0 μL of purified cDNA was amplified by PCR in a reaction prepared with 1.2 μL of 25 μM "UGA-V3" barcoded forward primer, 1.2 μL of 25 μM "UGB" reverse primer (primers listed in Table S7), 3.6 μL of water, 3 μL of 100% DMSO, and 15.0 μL of 2X Phire Hot Start II PCR Master Mix (Thermo Fisher #F125L). The following thermocycler protocol was used: 98 °C for 30 seconds; then 98 °C for 10 seconds, 66 °C for 10 seconds, and 72 °C for 10 seconds, repeated for 18 additional cycles; 72 °C for 5 minutes; and 4 °C hold. PCR products were purified with 2% E-gel Agarose EX gels cut and Qiagen MinElute PCR purification kits, with purified cDNA eluted in 25 μL of Qiagen EB buffer. The amplified DNA was quantified by Qubit HS dsDNA and sequenced on a UG100 Platform (Ultima Genomics, Fremont, CA).

Sequencing data were processed into SHAPE profiles with the Ultraplex-Bowtie-RNAframework (UBR) pipeline, available at https://github.com/daslab/ubr/. The SHAPE data were normalized so that the 90th percentile SHAPE value across all tested designs in a library was set to 1.0.

*FAST-MaP for design characterization with c-di-GMP*

To characterize ligand-dependent SHAPE responses, we used a faster turnaround protocol based on primerless sequencing called Fast and Accessible Sequencing Technology for Mutational Profiling (FAST-MaP). The W03 starting sequence and the top Eterna and MPNN-fixbb designs from Round 1, as well as the *V. cholerae* class I c-di-GMP riboswitch from ref. (*53*) (used as a positive control; PDB 3IWN), were synthesized, folded, and modified with 2A3 as described above for the main libraries, with three changes: the templates for each construct were ordered as gBlocks (IDT, Coralville, IA; see **Table S7**); each construct was prepared as two parallel reactions, with and without 5 mM c-di-GMP (Enzo, BLG-C057-01) added after $MgCl_2$ refolding, and incubated at room temperature for 20 min before modification; and final libraries were sequenced by Plasmidsaurus (Eugene, OR) with Premium PCR (6,000 reads per sample). The standard error on SHAPE reactivities was estimated from counting statistics by UBR or a recently accelerated data processing pipeline cmuts (https://github.com/hmblair/cmuts).

<u>OpenKnot Score</u>

The OpenKnot score builds on the Eterna score developed previously for non-pseudoknotted secondary structures (1). The Eterna score looked at all SHAPE-probed nucleotides, excluding flanking regions added to enable experimental characterization. If the nucleotide was asked to be paired in the target secondary structure but had SHAPE reactivity above 0.5, then it was considered unacceptable. If instead the nucleotide was asked to be unpaired in the target secondary structure, SHAPE reactivity below 0.125 was considered unacceptable. The Eterna score was defined as the percentage of target nucleotides considered acceptable given these thresholds. To emphasize proper achievement of pseudoknots, an additional Crossed Pair score was computed over just the nucleotides involved in pseudoknots in the target, i.e., $i$ or $j$ from target base pairs ($i$,$j$) such that there existed another target base pair ($k$,$l$) with $i < k < j < l$ or $k < i < l < j$. The OpenKnot score was defined as the average of the Eterna score and the Crossed Pair score. (In prior work, a version of this score called the ‘ensemble’ OpenKnot score was used to compare free-form pseudoknot designs to an ensemble of predicted structures; but here, the target secondary structure was pre-specified and we focused on match to the target secondary structure, previously called the ‘target’ OpenKnot score.) Only designs with estimated signal-to-noise ratios above 1.0 were considered for assessment and selection of designs for compensatory mutagenesis or cryo-EM. Scoring code is available at https://github.com/eternagame/OpenKnotScoreMATLAB and https://github.com/eternagame/OpenKnotScorePipeline. For target W09 in Rounds 1-2 and target P06 in Round 3, two structures were made available at different times to predictors (Tables S4 and S5); for each design, the OpenKnot score was computed with each of the two alternative structures and the design was assigned the maximum of these two scores. Experimental errors in OpenKnot scores were estimated based on the standard deviations across the 10 replicates of the starting sequence for each target; the average standard deviation was 4.9 (Round 1), 2.1 (Round 2), 1.1 (Round 3), and 1.9 (Round 4).

When comparing methods, an OpenKnot score greater than 90% was taken as a threshold for success; in figures, error bars on the fraction of ‘success’ targets reflect ±34.1% confidence intervals, analogous to standard deviations for normal distributions, here computed using the exact formula for the binomial distribution. OpenKnot scores were considered only for designs with sequence identity to the starting sequence lower than 80%. To remove low-quality data, the designs were further filtered for those whose SHAPE profiles had estimated signal-to-noise ratios greater than 1.0. The overall success rate per round for each method was defined as the number of targets for which a method succeeded divided by the total targets in that round (this included targets where no designs were submitted by some of the methods, such as in Rounds 1 and 2). In Rounds 3 and 4, to mitigate cases where design methods led to artefactually correct SHAPE profiles from distinct secondary structures, designs whose secondary structure predicted by RNet were different from the target secondary structure (with harmonic mean of precision and recall, $F_1 < 0.8$) were removed for final statistics. Statistical significance for differences between methods was based on paired tests using exact binomial distributions. In alternative analyses based on Z-scores, the 80th percentile OpenKnot score for each method on each target was used as a representative score, to correct for the different numbers of design submissions from different methods and to approximate a 'best of 5' evaluation scenario. Z-scores were computed as the number of standard deviations beyond the mean of these representative scores across all methods, with the mean and standard deviation recomputed after removal of any methods with $Z < -2$, to remove outliers. The sum of the non-negative Z-scores across targets were used to rank methods; ignoring Z-scores lower than zero prevented penalizing methods for idiosyncratic failures, as is carried out in CASP evaluations (8, 9). Significance in differences between summed Z-scores was estimated based on exact enumeration of the distribution of sums upon resampling.

Compensatory mutagenesis (M2R-seq)

For each target and each design method, the design with highest OpenKnot score in Round 3 was characterized by compensatory mutagenesis. As part of this selection, RNet-predicted secondary structures were computed, and designs with F1 accuracies over 0.8 (compared to the target secondary structure) were prioritized. If no designs exceeded this cutoff (as occurred for several Rosetta sets and for gRNAde and Struct2SeQ with P16), the top OpenKnot score design was selected; these served as negative controls (see, e.g., **Fig. S7**). For each design and for each target base pair, single mutants for each side of the pair and the compensatory double mutant were synthesized. The mutations converted each nucleotide to the sequence of its target pairing partner in the starting design, based on prior measurements (10, 11) as well as in silico studies suggesting that this choice would maximize the rescue signal. Flanking sequences, including unique barcodes, were added, and designs were characterized by SHAPE measurements as described above. The rescue factor was computed as follows:

$$Rescue\ factor = 1 - \frac{\sqrt{\sum_i \left(r_i^{AB} - r_i^{start}\right)^2}}{\sqrt{\sum_i \left(r_i^{A} - r_i^{start}\right)^2 + \sum_i \left(r_i^{B} - r_i^{start}\right)^2}},$$

where $r^{\mathrm{start}}$, $r^{\mathrm{A}}$, $r^{\mathrm{B}}$, and $r^{\mathrm{AB}}$ are, respectively, the SHAPE profiles for the starting design sequence, for the design mutated at one partner of the target pair, for the design mutated at the other target pair, and for the design mutated at both sites, and the index $i$ iterates over all positions in the target sequence. This expression returns values near 0 if the disruptions in SHAPE profile for the double mutant can be described as the independent sum of disruptions in SHAPE profiles from each single mutant, i.e., no 'rescue' observed upon compensatory mutagenesis. The expression returns values near 1 if the SHAPE profile for the compensatory double mutant is restored to match the SHAPE profiles of the starting sequence. Rescue factors over a threshold of 0.4 for any base pair in the stem were taken as evidence of stem formation, as calibrated in ref. (11) based on computational studies. Changing this threshold in the range 0.3-0.5 did not change the method rankings. The overall M2R-seq recovery score for a design was the percentage of target stems for which at least one base pair gave rescue upon compensatory mutagenesis.

Computational design methods

*3DRNA*

Designs were generated using 3DRNA, a deep neural network trained to perform sequence design on a fixed RNA backbone using atomic context. The method paralleled one of the early protein sequence-design protocols described in Anand et. al. (12). In brief, 3DRNA was trained on the crystal structures in the original training dataset described in gRNAde (13, 14). For each nucleotide, the model canonicalized along the C1′ atom, and identified the N, C, O, and P atoms in a 20 Å cube voxelized to a 0.5 Å resolution. A 3D convolutional neural network was then trained to classify the predicted base identity and binned $\chi$ angles from this atomic environment

Design calculations in Round 1 were aimed at designing RNAs with low sequence identity to wild type. For each OpenKnot problem, the 3DRNA protocol required a fixed backbone; therefore, each structure was prepared by rebuilding missing nucleotides if present (W06) or predicting structures if none were given (W09 - W13) with DeepFold RNA (15). Each backbone was subsequently minimized using Rosetta Relax with backbone constraints. To avoid initial bias, all starting sequences were randomized. Sequence design proceeded using 3DRNA following a blocked Gibbs sampling procedure using the Metropolis Hastings criterion, with Rosetta Repack after each proposed mutation was made. Each design trajectory consisted of 500 iterations. For each problem, 50 sequences were generated and evaluated for secondary structure consistency using EternaFold (16), and the top 20 candidates were selected.

This 3DRNA protocol was restricted to fixed-backbone 3D RNA design problems, and was used in Round 1 to explore sequence diversity under structural constraints. Due to limitations in RNA structure prediction and other constraints, we did not continue this structure-centric design approach in later Rounds. However, the 3DRNA protocol represents a structure-focused baseline.

Code and model weights for instruction and installation are available at: https://github.com/ProteinDesignLab/3DRNA

*gRNAde*

Designs were generated using gRNAde, a structure-conditioned autoregressive RNA language model. As detailed previously (14, 17), gRNAde represents a 3D RNA backbone structure as a geometric graph where each nucleotide is a node featurized via a 3-bead coarse-grained representation (P, C4', and N1/N9 atoms) and local geometric descriptors. An SE (3)-equivariant Graph Neural Network (GNN) encoder processes this geometric graph to produce latent representations, which are decoded by an autoregressive language model to predict the sequence from 5' to 3'.

For Rounds 1 and 2, an initial version of the model was employed as described in (14, 17). For Rounds 3 and 4, we utilized the final model architecture, which incorporated explicit secondary structure edge types into the graph connectivity, as described in (18). To enable robust design for targets lacking experimental 3D structures, the final model was trained with a specialized dropout strategy: for 50% of training examples per epoch, 3D backbone coordinates were masked, forcing the model to learn implicit 3D structural information solely from 1D sequence and 2D secondary structure connectivity. The final model was trained on a dataset of 4,211 RNA structures with up to 500 nucleotides from the PDB, covering a diverse set of geometries at resolution ≤ 4.0 A. This broad training distribution enabled the model to capture non-canonical interactions and tertiary motifs critical for stabilizing complex pseudoknots.

Rounds 1 and 2 served as ablation studies to calibrate computational filtering metrics and transition from manual to automated design. In both rounds, up to 1,000 candidate sequences were sampled per target using the initial model at a fixed temperature of 0.1. Designs were scored using four computational filters: 1D chemical mapping profile consistency using RNet (mean absolute error, MAE), secondary structure self-consistency using EternaFold (16) and RNet-SS (2) (Matthews correlation coefficient, MCC), and model confidence via gRNAde perplexity. While the pipeline initially evaluated 3D self-consistency using RhoFold (originally E2Efold-3d) (19), this metric was excluded after control studies revealed that the predictor failed to recover the correct backbone geometry for native sequences. Given this inability to reliably fold the 'ground truth' inputs, we concluded that current 3D prediction tools are not sufficiently robust to serve as filters for de novo designs.

In Round 1, selection relied on human-in-the-loop inspection. The submissions were a mix of unconstrained global designs and designs subject to manual constraints, such as fixing nucleotides at stem-loop termini or within pseudoknots. Retrospective analysis revealed that manual selection was unscalable and that unconstrained designs performed comparably to constrained ones. Crucially, we observed that RNet-MAE correlated strongly with experimental success, while the secondary structure MCC scores showed moderate correlation and perplexity showed poor correlation.

In Round 2, the pipeline was automated to rigorously stress-test these findings. Constraints were removed, and the top 5 designs were selected automatically for each scoring metric (RNet MAE, EternaFold MCC, RNet-SS MCC, and gRNAde perplexity), resulting in 20 total designs. Because this round was treated as a controlled ablation study, including candidates ranked by weaker metrics like perplexity rather than exclusively optimizing for the strongest predictors, the aggregate performance of gRNAde was lower relative to methods that fully exploited the best available filters. However, this unbiased comparison was essential to confirm that 1D chemical mapping predictions from RNet provided the most reliable signal for prospective design filtering.

Based on the calibration in earlier rounds, a fully automated, high-throughput pipeline was deployed for Rounds 3 and 4. The pipeline utilized the final gRNAde model and a highly parallelized implementation of the RNet scoring system, enabling the screening of 1 million designs per target in under 12 hours on a single NVIDIA A100 GPU. This high-throughput capability represented a significant advance; while physics-based methods like Rosetta required hours of CPU time per design, gRNAde generates hundreds of designs in approximately 1 second on a GPU. This speed enabled the exploration of a sequence space orders of magnitude larger than previously possible.

To maximize diversity, sequences were sampled across a varying temperature range of 0.1 to 1.0. This range allowed the model to balance high-confidence, lower-diversity sampling (low temperature) with broader exploration of the solution space (high temperature). Replacing the generic MAE metric in previous rounds, computational filtering was refined to use an in silico OpenKnot score simulated via RNet, which was also shared with all competitors (https://github.com/chaitjo/geometric-rna-design/blob/main/src/openknot_score.py). Designs for natural targets were first filtered for secondary structure consistency (MCC > 0.9 via RNet-SS). Crucially, secondary structure filtering for synthetic targets was not applied, as RNet-SS was fine-tuned on natural sequences and was hypothesized to not generalize to non-natural topologies. All designs were then ranked by the in silico OpenKnot score to select the top 20 candidates.

For these rounds, we submitted two distinct sets of 20 designs per target, generated by the same model weights to assess the necessity of explicit 3D inputs: (1) designs conditioned on both the target 3D backbone and pseudoknotted secondary structure, and (2) designs conditioned solely on the secondary structure (called ‘gRNAde-no3d’). For synthetic targets lacking experimental structures, we utilized predicted backbones provided by the organizers and other participants (generated via trRosettaRNA, FARFAR2, or RFDpoly) for the 3D-conditioned set. This ablation demonstrated gRNAde's capability to implicitly model 3D constraints even when explicit backbone coordinates were unavailable.

Code: https://github.com/chaitjo/geometric-rna-design

*NA-MPNN and RFDpoly*

Three methods were tested using NA-MPNN for sequence design and RFDpoly for structure generation: MPNN-RFdiff, MPNN-fixbb, and codesign-RFdiff. In the MPNN-RFdiff approach, secondary structures for each puzzle were provided to RFDpoly (20) in dot-bracket notation; these secondary structures were used to construct 2D base pair templates, which conditioned structure-denoising trajectories to generate RNA backbones accommodating the specified base pair networks. In all rounds, the resulting diffusion-generated coordinates were then given as inputs to NA-MPNN (21), which autoregressively predicted RNA sequences for the given backbone. For Rounds 3 and 4, the RFDpoly-generated models produced by conditioning on each target secondary structure were made available to other competition participants, to support use of 3D structural information in alternative sequence design approaches. To disentangle performance of sequence design from structure generation steps, we also explored an approach denoted MPNN-fixbb, wherein NA-MPNN sequence design was performed on alternative backbone models: experimentally derived structures when available, or alternative (non-RFDpoly-generated) computational models otherwise. An additional method, codesign-RFdiff, involved sequence-structure co-generation, in which random positions had their predicted sequence tokens revealed autoregressively over the last 40 steps of a 50-step structure-denoising trajectory, utilizing RFDpoly's ability to predict sequence in addition to structure during each forward pass. In Round 1, both structure generation and sequence design were performed using preliminary versions of RFDpoly and NA-MPNN from early stages of model development, whereas subsequent rounds used the final model versions (available from the GitHub pages linked below), as reported in their respective manuscripts. For all three methods, final submitted sequences were filtered based on in silico predictions of secondary structure, given designed sequences. For Round 1, designed sequences were filtered for predicted $F_1$ score self-consistency using EternaFold (16) to predict secondary structures, which were compared to the target secondary structure for each puzzle. In OpenKnot Rounds 2 through 4, RNet (2) predictions were used instead: for designed sequences and their corresponding puzzle's target secondary structure, RNet 1D predictions were used to compute predicted Target OpenKnot scores, and RNet-SS 2D predictions were used to compute predicted $F_1$ scores. Sequences that passed RNet filtering were then further filtered for self-consistency by predicting tertiary structures (using Chai-1 (22) for Round 2 and AlphaFold 3 (23) for Rounds 3 and 4), from which base pairs were extracted and again used to compute predicted $F_1$ scores.

Code and instructions for installation and inference can be found at the following links.
RFDpoly: https://github.com/RosettaCommons/RFDpoly
NA-MPNN: https://github.com/baker-laboratory/NA-MPNN

*Rosetta*

Rosetta RNA designs were created with the rna_design executable (2016 build; rosetta_bin_linux_2016.32.58837_bundle) available in the Rosetta3 codebase. This algorithm carries out simulated annealing on side chain conformations and identity given an input RNA backbone, using either a default high resolution score function ('rna_hires.wts') or a low resolution score function (‘rna_lores.wts’), both originally hand-crafted for RNA 3D structure prediction (24). For Round 1, 20 submissions were made for each score function, using the 3D structures supplied by challenge organizers. For Round 2, no further submissions were made. For Round 3, the Rosetta runs were scaled up to 5000 designs each for the high resolution and high resolution score functions followed by RNet filtering, as in gRNAde. The 20 submissions were based on the 10 with best RNet scores and then another set of 10 using lowest sequence identity. For Round 4, targets were too large for effective computational design.

Code: https://github.com/RosettaCommons/rosetta

*Struct2SeQ and Struct2SeQ-SHAPE*

Struct2SeQ is a reinforcement learning framework that directly learns to generate RNA sequences from target secondary structures. By leveraging deep Q-learning and dense, chemically informed rewards derived from neural predictors of folding and chemical reactivity, the model enabled structure-conditioned generation of RNA sequences that conform to both structural and chemical constraints (2, 25). Struct2SeQ used two versions of RNet as the environment that provided reward signals to the reinforcement learning agent. RNet is a deep neural network designed for RNA structure-function prediction, capable of mapping a given RNA sequence to a SHAPE reactivity profile or, in a fine-tuned version called RNet-SS, to a base-pairing pattern. The structure-predictive RNet-SS enabled position-level rewards for correct base-pairing, while the SHAPE-predictive RNet enabled constraint-based filtering using thresholds on predicted reactivities.

The reinforcement learning model in Struct2SeQ is a neural network that involves an embedding layer that encodes the dot bracket input, sinusoidal positional encoding, 1D convolutions, graph convolutions using secondary structure connectivity, a transformer encoder and a transformer decoder. The model was trained with deep Q learning. In this setting, the model acts as an agent that incrementally constructs a sequence, making a discrete decision (nucleotide selection) at each position. After generating a full sequence, the agent is updated with scalar rewards on each position based on if that position is correctly paired or unpaired and if that position meets SHAPE constraints as defined by OpenKnot score. Two versions of Struct2SeQ were trained, one with only secondary structure reward (Struct2SeQ), and one with secondary structure and SHAPE reward (Struct2SeQ-SHAPE).

During inference, Struct2SeQ acts as an autoregressive RNA generator conditioned on a target secondary structure, producing Q-values for each nucleotide choice and using them to guide decoding. Sequences are generated one base at a time, with options such as beam search for high-quality, reliable folding, or stochastic sampling for increased diversity. A rescue strategy locally repairs near-miss sequences by enumerating corrections at mismatched positions, while post-hoc screening metrics—such as base-pair Jaccard similarity and OpenKnot scores—filtered generated candidates to ensure structural accuracy and SHAPE-compatibility. Designs for both Struct2SeQ models were submitted for the 100-mer targets in Round 3. Additional training time was required for achieving designs with satisfactory in silico scores for the longer 240-mer targets; as a result, Struct2SeQ designs for Round 4 were not prepared and submitted for experimental characterization until after release of experimental results for other design methods in Round 4.

Code: https://github.com/Shujun-He/Struct2SeQ

Cryo-electron microscopy

RNA designs for the P20 target ('Kissing Multiloops') were prepared for cryo-electron microscopy as follows, with full sequences given in Table S8. For each P20 design, the corresponding sequence was added to a circularly permuted group II intron scaffold to facilitate structure determination via cryo-EM. Each P20 construct was split at the tetraloop of the P6 stem for scaffolding. For the gRNAde and Struct2SeQ designs, a G was prepended to the design to aid transcription by T7 RNA polymerase. Genes for the P20 scaffolded constructs were ordered through GeneWiz and subsequently transformed into DH5α cells. Purified plasmid was linearized using an engineered BamHI reconstruction site at the end of the intended construct. RNA was in vitro transcribed by adding 50 µg of template plasmid to a total volume of 1 mL of in vitro transcription buffer (50 mM Tris-HCl pH 7.5, 25 mM $MgCl_2$, 5 mM DTT, 2 mM spermidine, 0.05% Triton X-100, and 5 mM of each NTP). T7 RNA polymerase and thermophilic inorganic phosphatase was added to begin RNA synthesis. The reaction mixture was incubated at 37 °C for 3 hrs. CaCl2 was added to a final concentration of 1.2 mM along with 10 µL of Turbo DNase and placed at 37°C for 1 hour to fully digest the DNA template. Proteinase K was subsequently added and incubated at 37°C for an additional hour to degrade the various protein components in the reaction mixture. The in vitro transcription reactions were then run over a size exclusion column (HiLoad® 16/600 Superdex® 200 pg) in a running buffer of 4 mM $MgCl_2$ and 5 mM Na-cacodylate pH 6.5. Elution fractions containing peaks that corresponded to well folded RNA scaffolds were collected. The RNA-containing solutions were then mildly refolded by increasing the $MgCl_2$ concentration to 10 mM and placing the solution at 50°C for 30 minutes. The RNA was then allowed to cool for 30 minutes at room temperature and then concentrated using a 100 kDa molecular weight cut-off filter to ~5 mg/mL. All cryo-EM grids were prepared by applying 3.5 µL of the appropriate RNA to a glow discharged (40 mBar, 15 mA for 40 s using a PELCO easiGlow) Au R 2/1 200-mesh grid (QUANTIFOIL®). The grids were blotted with filter paper (Whatman No.1) at 4°C in a cold room before plunge freezing into liquid ethane/propane (37.5/62.5) mix using a manual plunger.

To minimize potential experimental bias, cryo-electron microscopy of RNA was carried out without knowledge of the design sequences. Below, we describe a consensus protocol used to process cryo-EM data. Detailed processing workflows for individual RNA designs are provided in **Fig. S8** to **Fig. S11**, and special considerations for the Mol11 (MPNN-RFdiff) dimer given below.

Images were collected on Janelia Krios4 equipped with a cold field emission gun (CFEG), Selectris X energy filter and Falcon 4i camera. EER movies were imported and processed in cryoSPARC v4.5.3 (26). Motion correction and contrast transfer function estimation were performed using cryoSPARC's default settings. Initial particles were automatically picked using 2D templates generated from a reference volume of the group II intron scaffold (EMD 45247 (27)). Particles were extracted at a box size of 320px or 420px and binned by a factor of 2. The initial particle stacks were subjected to ab initio reconstruction into four to six classes. Typically, one or two classes corresponding to the group II intron scaffold with the circularly permuted RNA design were identified, and the particles were sorted by one or two rounds of heterogeneous refinement. In all cases, the RNA design showed no interactions with the group II intron scaffold.

A consensus reconstruction was created by non-uniform refinement or local refinement on the scaffold. Subsequently, the particles were reextracted to 420px at full pixel size or binned by a factor of 2. Density belonging to the RNA design (the sub-particle) was inspected by 3D classification. 3D classes with well folded RNA sub-particles were selected for further processing. Reference-based motion correction was performed using the final particle stack and volume as inputs. To align sub-particles, the density of the group II intron scaffold was removed by two successive rounds of particle subtraction. The remaining sub-particles were realigned using local refinement. Map sharpening was performed in cryoSPARC or PHENIX (28). Atomic models of the RNA designs were built de novo and refined in Coot (29) and PHENIX. For ease of viewing, residues have been renumbered to match the design target numbering for this manuscript; PDB depositions include numbering appropriate for the full length sequence, including the inserted group II intron scaffold.

For the Mol11 (MPNN-RFdiff) dimer, an atomic model for two complete monomers could be built into the dimer density, but some additional density was visible at the helical dimer interface (Movie S2), which appeared due to the transcription of a remnant of the BamHI restriction site (GGAUC) at the 3′ end. Base pairing between the BamHI-derived sequences from each monomer was modeled as a paired region ('P8') and constituted one of three dimer interfaces, together with pseudoknot P4 and base stacking of A28 from the GAAA loop of P3. The BamHI sequence was not visible in any of the other maps.

<u>3D structural comparisons</u>

Secondary structures and lists of canonical and non-canonical base pairs were extracted from 3D structures with DSSR (30). TM-align comparisons of 3D structures were carried out with the US-align package (31), using the C1′ atom as the representative atom for each nucleotide. Identification of the most similar previous 3D structures was carried out on a directory containing all RNA chains and entries in the PDB as of December 17, 2025. Backbone-independent structure search was carried out with ARTEM 2.0 (32) against the database 'RNA_nrlist_3.363_3.0A', available through webserver at https://genesilico.pl/artemws/motif_search. The tertiary interaction from the MPNN-fixbb design was isolated as the nine residues 1, 57-58, 76-77, 90-93, and the ARTEM search was carried out requiring that all residues find a match. The tertiary interaction from the gRNAde design was isolated as the 12 residues 36, 63-65, 74-75, and 81-86; ARTEM searches forcing all 12 residues to match failed to find matches, so partial matches with as few as 10 residues were tabulated.

<u>Use of AI-assisted technologies</u>

Anthropic's Claude (Opus 4.6 and Opus 4.7) was used during preparation of the revised manuscript. Specifically, Claude assisted with prose editing, drafting and refining figure captions, drafting components of the response to reviewers, and auditing the internal consistency of cross-references between the main text, supplement, figures, and tables; and Claude Code was used to write and execute Python code for computational analyses underlying **Fig. S2**, **Fig. S6,** and **Fig. S14**. All AI-generated text and analyses were reviewed, edited, and verified by authors before incorporation. The authors take full responsibility for the accuracy of all content.

### Supplementary Text

<u>Ligand binding to de novo designs</u>

The W03 target is the secondary structure of the cyclic-di-GMP-II riboswitch from *Clostridium acetobutylicum* (PDB 3Q3Z(*25*)), which is stabilized into the pseudoknot conformation by binding of the c-di-GMP ligand. Throughout the OpenKnot challenge, designs were probed in the absence of small molecule ligands so that the formation of the target secondary structure would reflect the de novo design itself rather than ligand-induced stabilization of an otherwise unstable structure. To examine whether this experimental choice obscured differences between designs that did or did not retain ligand-dependent function, we re-probed the W03 starting sequence and the top Eterna and MPNN-fixbb designs from Round 1 (**Fig. 1A-C**) in the presence and absence of c-di-GMP with SHAPE (**Fig. S2A-B**).

Both the Eterna and MPNN-fixbb designs achieved high OpenKnot scores both with and without c-di-GMP, with no substantial change in OpenKnot score upon ligand addition **(Fig. S2**). This insensitivity is expected: the OpenKnot score is computed from SHAPE reactivities and is therefore mainly sensitive to base pairing, which does not change appreciably upon ligand binding for these designs; the pseudoknot is already pre-organized in solution.

Despite this insensitivity at the level of secondary structure, the Eterna design exhibited reproducible local SHAPE protections upon c-di-GMP binding at nucleotides corresponding to the ligand-binding pocket of the natural riboswitch (**Fig. S2E**). This pattern is consistent with the Eterna design having retained the noncanonical contacts that recognize c-di-GMP, an outcome enabled by the design's conservative re-design strategy that preserved key putative ligand-contacting residues while diversifying the rest of the sequence. By contrast, the MPNN-fixbb sequence, a more aggressive de novo design with low sequence identity to the natural riboswitch (<50%; **Fig. 1F**), showed no localized protection upon c-di-GMP addition, consistent with loss of ligand recognition. These results confirmed that pseudoknot secondary-structure design and the preservation of small-molecule recognition are separable design objectives that can each be addressed independently.

As a positive control for the SHAPE-based detection of ligand binding, we probed a class I c-di-GMP riboswitch from Vibrio cholerae (PDB 3IWN (*53*)), which forms its pseudoknot only upon c-di-GMP binding. The class I riboswitch displayed clear ligand-induced protections at nucleotides corresponding to the canonical c-di-GMP contacts in the deposited 3D structure, confirming that our SHAPE assay can detect ligand-induced local protection when it occurs (**Fig. S2F**). Unexpectedly, under our buffer conditions, the natural W03 starting sequence (the c-di-GMP-II class natural sequence) did not form its target pseudoknot even at saturating c-di-GMP (5 mM), in contrast to the originally published behavior (**Fig. S2A**). We attribute this to differences in our construct, which lacks a 5′ guanosine present in some published constructs and includes flanking sequences that may interact with the aptamer. We did not pursue this discrepancy further, as the focus of this work is on the designability of pseudoknot secondary structures in the absence of stabilizing partners.

<u>Player approaches to design</u>

Eterna participants responsible for these high-scoring designs used a combination of in-game tools, external software, and design strategies. This section summarizes design methods used by four of the participants with consistently high performance across Eterna challenges.

*Ucad*

The vast majority of my designs were based on specificity rules. For each designed sequence, I'd select a pattern and use it for every loop. Usually all A or all U with a C to avoid excessive repeats in longer loops. Next, I'd create a diverse set of stem sequences while avoiding including the complement of the loop. (When using A loops, I'd often ban UpU from stem sequences.) Intended to reduce misfolding and ensemble diversity.

Some structures had known or predicted pairings provided to the players, particularly in round 1. I tried to conserve those non-canonical pairings or triple helices.

*Eli Fisker*

In the case where we had PDB files, I looked at the structures in Chimera and used that to try to lengthen the triplex area (if present) or create one. (Not sure I was successful) In puzzles with riboswitches that didn't have their binding partner (like fluoride for PreQ1-II), I mutated in the area where the interaction would have been, trying to make a binding internally instead.

I created a series of 5 different designs in some of the R3 puzzles. The goal was to stabilize a solution in RNet with as high an F1 score for target structure and pseudoknot as possible, while at the same time maximizing for other characteristics:

- Maximum amount of GC's possible
- Maximum amount of AU's possible
- Maximum amount of GU's possible
- Maximum amount of base mutations
- Maximum amount of cross-pair score

My aim was to see if I could make great F1 scores in silico, while provoking failure in vitro, testing the limits of the RNet algorithm, because I know from earlier lab experience that some of the above design types can fail.

In puzzles where a structure had SHAPE data from an earlier round, I identified bases in the structure where SHAPE suggested it could be improved. I ran mutation series on them with Jeff's ArcKnot tool, maximizing the F1 cross-pair score.

In the puzzles that came from natural origin sequences, I ran a BLAST and looked at what mutations were common in the MSA viewer. For the novel structures, I ran mutation series on a design I had optimized for F1.

*mjt*

I relied heavily on the parameters such as low as possible amount adenines, as high as possible confidence values for target mode and pseudoknots, and trying to achieve or get as close as possible to target mode. That is all that I did, just try to improve the parameters as much as possible. I do think playing the games and the labs helped me get a feel for picking what bases to try to get the responses I wanted.

*Jandersonlee*

For round 4, I used my ArcKnot booster tool that runs within the Eterna lab tool. This booster provides visualization (as a dot-plot or arc-plot) for the base pairing probability data as computed according to RNet as well as computing the F1 and F1 cross-pair metrics that evaluate, among other things, the prediction of the presence of a pseudoknot in the modeled sequence secondary structure. This tool can also be used to perform multiple rounds of single-base or base-pair modifications on a design in an attempt to improve upon the metrics.

I began by reviewing the designs submitted by other players and selecting ones that seemed promising in terms of their dot-plots and metrics as having some evidence of a plausible pseudoknot forming according to the selected energy model. I then selected a few designs for each puzzle and used them as starting points for one or more mutation rounds, again using the ArcKnot tool. Finally, I ranked the designs according to F1 score or F1 cross-pair score, and submitted 5 to 10 of the top designs for each of the starter sequences.

Thus, my selection of sequences for submission was based on a combination of experience-mediated inspection, and assessment of starting sequences and their “mutated” counterparts aided by energy-model based dot-plots, design metrics, and metric-mediated mutation rounds, using a browser-based tool that I had created.

Round 1 performance gap between AI and Eterna

Despite success in the target W03, neither MPNN-fixbb nor any other AI method achieved OpenKnot scores > 90 in half of targets, worse than the 9/17 (47%) performance achieved by simply re-using the starting sequence provided with each target (**Fig. 1H**). When gRNAde and MPNN-fixbb were evaluated solely on the targets with PDB structures, they achieved OpenKnot scores > 90 at a higher rate, but still with just under 60% of targets (light bars, **Fig. 1H**). Even when grouped together, all the AI methods gave significantly lower performance than Eterna participants, with OpenKnot scores >90 in 11/17 (69%) vs. 16/17 (94%) targets, respectively ($p = 0.031$; exact paired binomial test on discordant pairs, one-sided).

Secondary-structure filtering of Round 3 and 4 designs

In Round 3, only one target was not designable with OpenKnot score > 90 by any method or Eterna participant (P16, "AK_PK100-3"; **Fig. 2B**). Indeed, while some AI designs submitted for this RNA received high OpenKnot scores from SHAPE, RNet predicted that these designs formed alternative secondary structures that fortuitously showed SHAPE profiles similar to that expected for the target. To be conservative, these designs, some of whose stems were confirmed in subsequent M2 and M2R experiments to be incorrect (see **Figs. S6** and **S7**), were filtered out in assessing performance in Rounds 3 and 4 (solid bars vs. light colored bars in **Figs. 2C** and **2F**).

Detailed analysis of cryo-EM structures

All three designs imaged as monomers by cryo-EM shared a similar global tertiary fold and coaxial stacking pattern (**Fig. 4C-E** and **Movie S1**); the TM-align scores of the MPNN-fixbb and gRNAde designs to the Struct2SeQ-SHAPE design were 0.511 and 0.535, respectively, both larger than 0.45, the cutoff for a matching global fold (*54*). Furthermore, all three global folds were distinct from the fold predicted by AlphaFold 3 (compare **Fig. 4B** to **Fig. 4C-E**; TM-align values: Struct2SeQ-SHAPE, 0.298; MPNN-fixbb, 0.304; and gRNAde, 0.250, all lower than 0.45). As expected, given their derivation from a human-imagined pseudoknot secondary structure, the designs were not structurally homologous to any RNA 3D structures previously deposited in the PDB (highest TM-align to prior PDB structure of 0.40, 0.35, and 0.38 for Struct2SeQ-SHAPE, MPNN-fixbb, and gRNAde designs, all lower than 0.45).

Detailed inspection of the two AI designs imaged at highest resolution revealed additional surprises in the form of intricate non-canonical interactions. In the MPNN-fixbb design, the 5′ terminal nucleotide G1, which should have base paired within the P1 helix in the target secondary structure, instead split off of P1 to form a non-canonical base pair with A92 in J5/7 (**Fig. 4D**), an interaction we term a ‘5′G/receptor’. In addition, the first two base pairs of P3 were rearranged to produce a register-shifted stem with an extrahelical bulge and an undesigned noncanonical base pair with the J4/1 junction (C34-A41). In the gRNAde design, nucleotides at two joining regions (J4/1 and J5/7) formed Watson-Crick-Franklin pairs that were not part of the P20 target secondary structure, ‘gluing’ together the adjoining stems into coaxial stacks (**Fig. 4E**). Remarkably, the tetraloop L7 did not form the compact loop expected for this GAAA sequence. This loop instead flipped out its last nucleotide A85, which intercalated into a sequence-distal receptor composed of P4 and P6 stems. This 'bulged-A/receptor' interaction involved a noncanonical pair of A85 with nucleotide G64 (**Fig. 4E**), which was itself therefore not available to base pair with its target partner C75 at the edge of P6. This interaction was further stabilized by stacking with a base triple formed between the excluded P6 nucleotide C75 and the G36-C63 base pair within the P4 helix.

In principle, the non-canonical base pairs, base triplets, and tertiary interactions visualized by cryo-EM (**Fig. 4D-E**) could have arisen due to learning and explicit design of these interactions by the gRNAde or MPNN neural networks, but several lines of evidence disfavor this picture. First, while both networks were originally developed with filters based on 3D predictions of AlphaFold-like models, those sequence-to-structure models do not predict these non-canonical features and, as noted above, mispredict the global fold; furthermore, gRNAde designs did not use 3D prediction as filters in this study. Second, all the cryo-EM noncanonical interactions involved breaking of target canonical pairs of the P20 secondary structure, despite the primary design criterion being the formation of these target pairs. Third, while the RNet neural network, used to filter the AI designs, could potentially have learned these non-canonical features, RNet predictions for these sequences' secondary structures recovered the target secondary structures including the P1 and P3 edge pairs that were broken in the MPNN-fixbb design cryo-EM structure. Fourth, while both gRNAde and MPNN-fixbb were guided by input 3D backbone structures (modeled, in this case, by RFDpoly and FARFAR2, respectively), visual inspection and the DSSR tool (*55*) confirmed that while most input models showed close approach of P6 and P7 helices and a few showed A85 bulging out of L7, none displayed any of the cryo-EM-observed noncanonical pairs, triplets, or tertiary interactions. Fifth, both gRNAde and MPNN were trained on the full database of prior RNA 3D structures, which could have contributed templates for the interactions, but we did not find clear templates using Aligning Rna TErtiary Motifs (ARTEM) (*56*). For the 5´G/receptor interaction in the MPNN-fixbb design, ARTEM found homologs in ribosome structures and a streptomycin aptamer (PDB: 1NTA (*57*)), but with pyrimidines substituting for the 5´G in each case (**Fig. S13A**). For the bulged-A/receptor in the gRNAde design, the best ARTEM matches were incomplete, visually poor, and did not recover the bulged-A flipping out of an apical loop distal from the receptor (**Fig. S13B**). These analyses suggest that the noncanonical tertiary features observed in these cryo-EM structures did not occur through explicit modeling by the AI methods. Instead, these features may have arisen through an inherent tendency of RNA nucleotides at the edges of helices to find arrangements that maximize non-canonical interactions. Interestingly, target base pairs whose nucleotides participate in such noncanonical interactions showed low M2R-seq rescue factors, suggesting that compensatory mutagenesis data, if scaled to base pairs for millions of designs, could inform future versions of RNet.

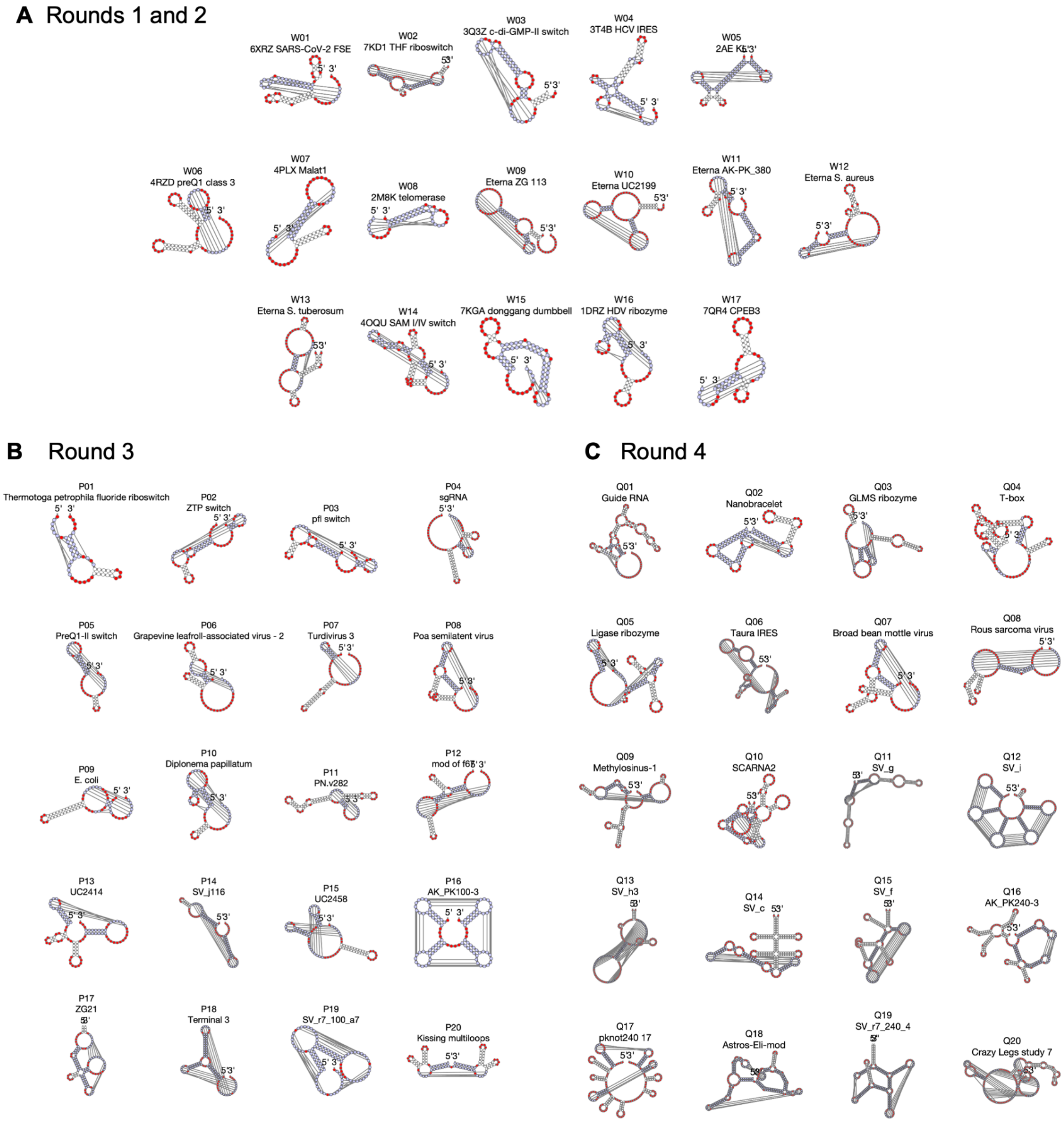

**Fig. S1. Target pseudoknot secondary structures for four rounds of the OpenKnot AI challenge.** Coloring shows target unpaired residues (red) and target paired residues (white or light blue, with the latter marking crossed pairs involved in pseudoknots).

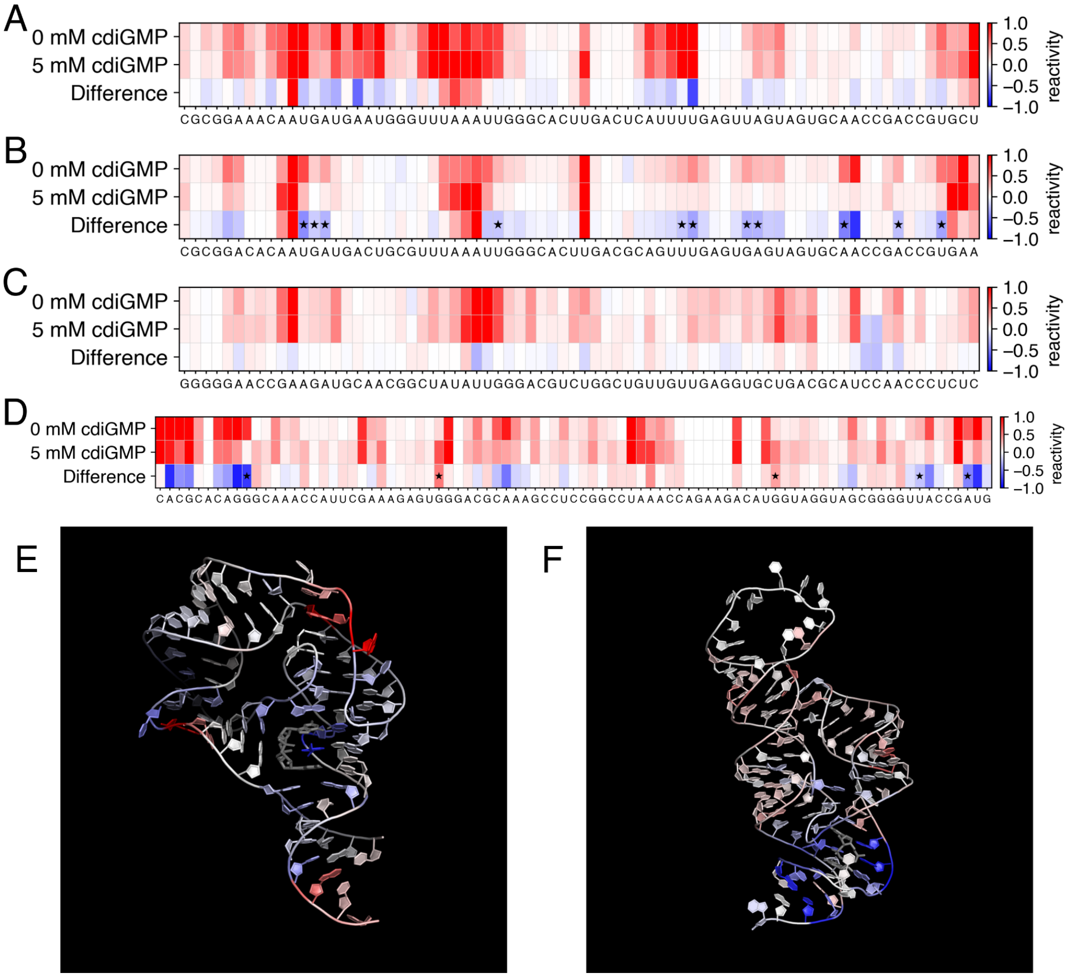

**Fig. S2. Pseudoknot secondary structure design and small molecule recognition are separable objectives in W03 (c-di-GMP-II riboswitch) designs. (A–C)** SHAPE reactivity at 0 mM and 5 mM c-di-GMP for the W03 starting sequence **(A)**, top Eterna design **(B)**, and top MPNN-fixbb design **(C)** from Round 1; rows show reactivity without and with ligand and their difference (blue, protected; red, increased exposure). OpenKnot scores were within error (~5) across conditions [70.1/74.1 (starting sequence); 91.3/91.2 (Eterna); 92.1/86.8 (MPNN-fixbb)] confirming pseudoknot formation independent of ligand. The Eterna design did not change c-di-GMP binding loops compared to the starting sequence and shows localized protections at the c-di-GMP binding pocket, consistent with ligand recognition; the *de novo* MPNN-fixbb design shows none. **(D)** Same analysis for a class I (3IWN, *V. cholerae*) riboswitch positive control, which folds only on ligand binding. Asterisks in (**A-D**) mark positions where the difference exceeds 3 times the standard error and mean SHAPE reactivity < 0.5. **(E,F)** SHAPE differences from (B,D) mapped onto the class II (3Q3Z) and class I (3IWN) crystal structures; protections (blue) localize at the c-di-GMP binding site (gray sticks). The WT class II construct (**A**) did not fold under our buffer conditions even at 5 mM c-di-GMP; see Supplementary Text.

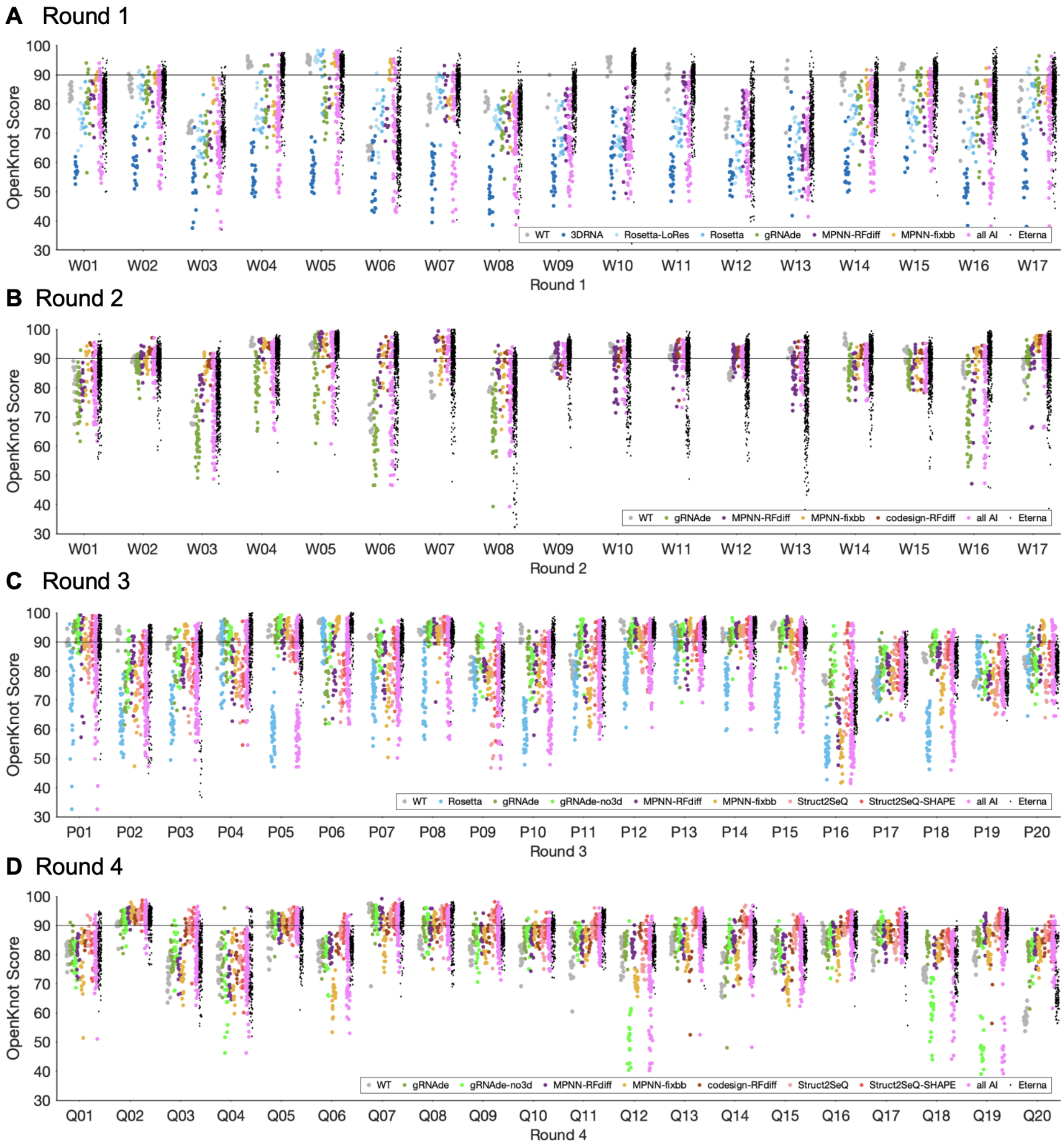

**Fig. S3. Performance for four rounds of the OpenKnot AI challenge.** OpenKnot scores for **(A)** Round 1, **(B)** Round 2, **(C)** Round 3, and **(D)** Round 4, separated by design method.

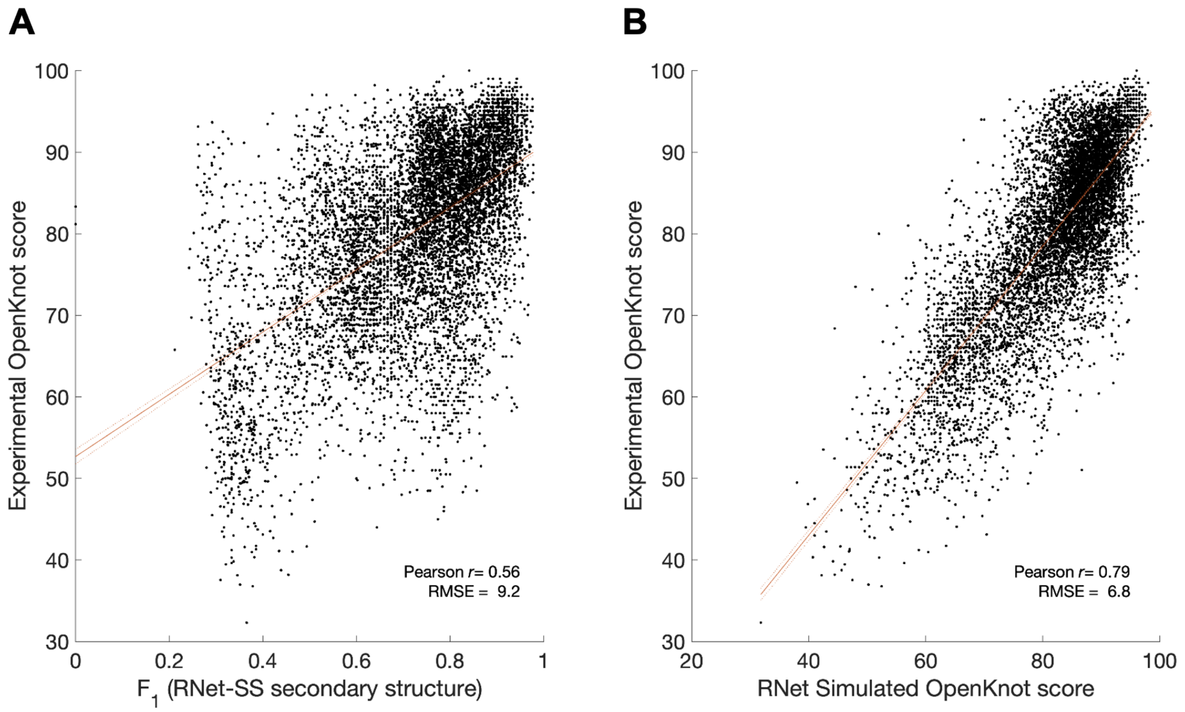

**Fig. S4. In silico metrics compared to experimental OpenKnot scores for Round 1.** In each panel, experimentally measured OpenKnot scores are compared to (**A**) accuracy of RNet-SS-predicted secondary structure compared to target secondary structure, as assessed by harmonic mean of precision and recall of base pairs $F_1$ and (**B**) simulated OpenKnot score using RNet-predicted SHAPE profiles.

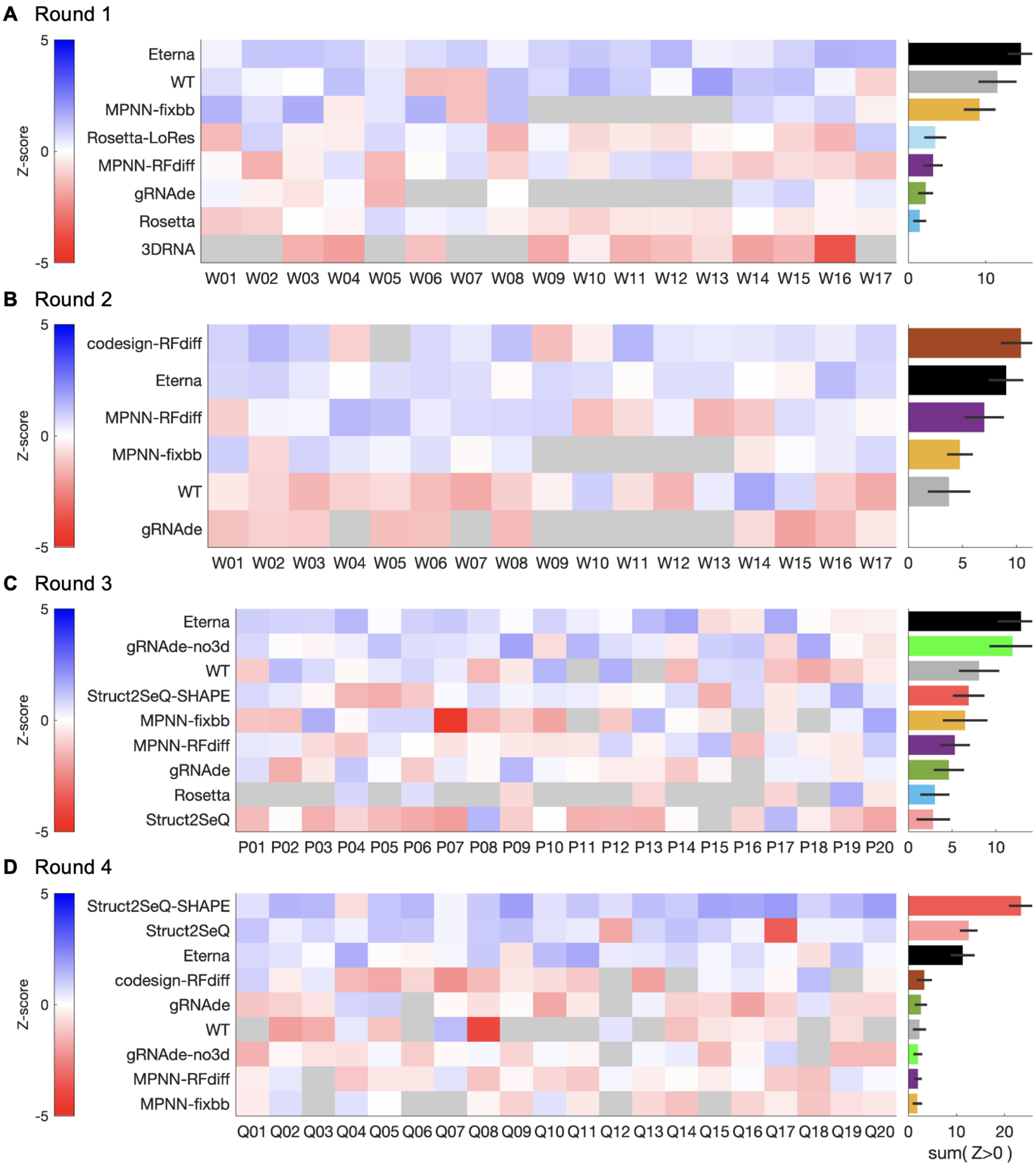

**Fig. S5. Z-score assessment for four rounds of the OpenKnot AI challenge.** To account for the different numbers of designs from different methods, the 80th percentile OpenKnot scores were tabulated for each target. The Z-score is the number of standard deviations over these scores, averaged across methods, after removal of outliers (Z<–2). Methods are ranked by the sum of their Z-scores, with negative Z-scores clipped to zero. The ‘all AI’ and other grouped methods are not shown due to poor performing subsets in these groups generally reducing their Z-scores.

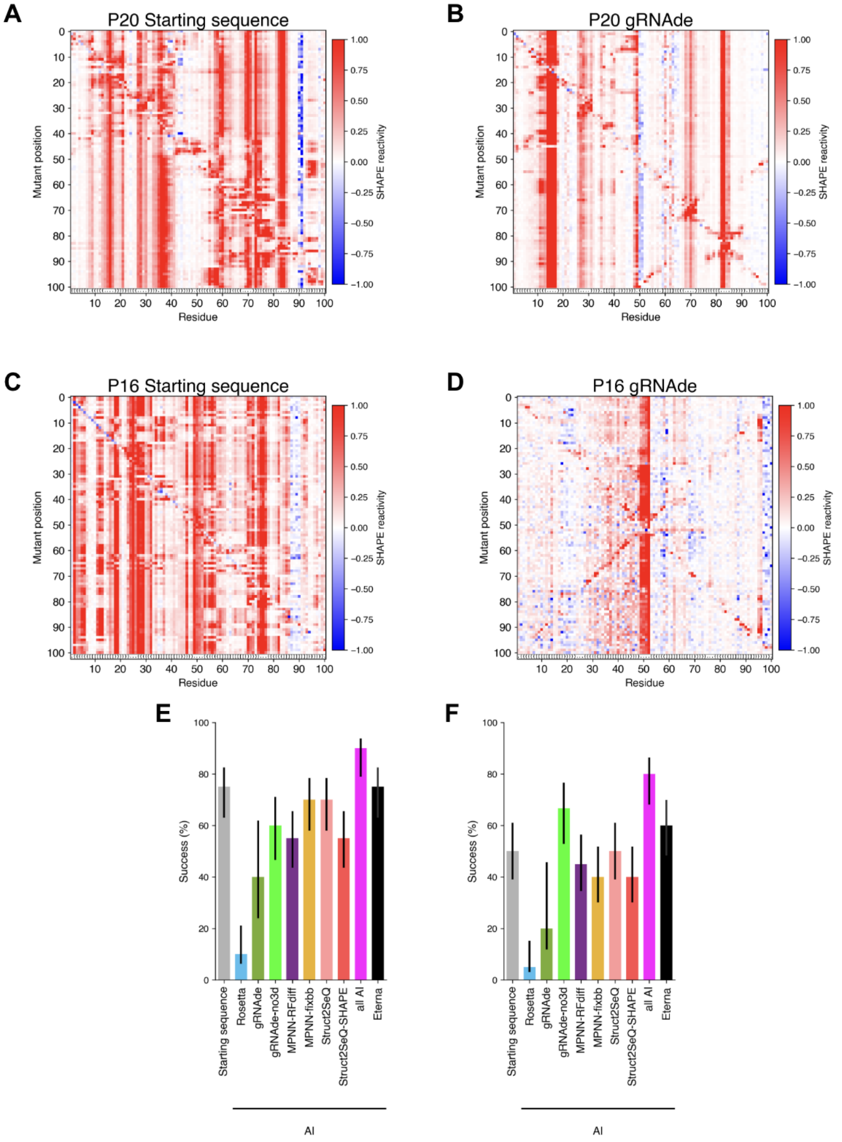

**Fig. S6. Two-dimensional mutate-and-map (M2) chemical probing and automated structure inference of OpenKnot Round 3 designs.** Two-dimensional mutate-and-map (M2) chemical probing (*34*, *35*) generates a SHAPE reactivity profile for the wild-type sequence and for single-nucleotide mutants of every position to its complement. **(A–D)** M2 data for P20 ('Kissing Multiloops'; **A,B**) and P16 ('AK_PK100-3'; **C,D**), for the target's starting sequence (**A,C**) or the gRNAde design **(B,D)**. The P20 designs (**A,B**) display clear correlated reactivity changes between distant positions confirming base-pairing contacts; not all mutations lead to precise release of their partners due to residual stacking or stabilization of alternative states. M2 data for the P16 starting sequence (**C**) does not show evidence of stable stems, while M2 data for the gRNAde design (**D**) shows an incorrect long hairpin (see also **Fig. S7**). **(E)** M2-guided inference of secondary structures with ShapeKnots suggests success ($F_1 \geq 0.8$) across the majority of 20 Round 3 targets by Eterna and AI methods, though this result is possibly biased by the energetic preferences of ShapeKnots, whose data-free predictions for these sequences produce similar results (**F**).

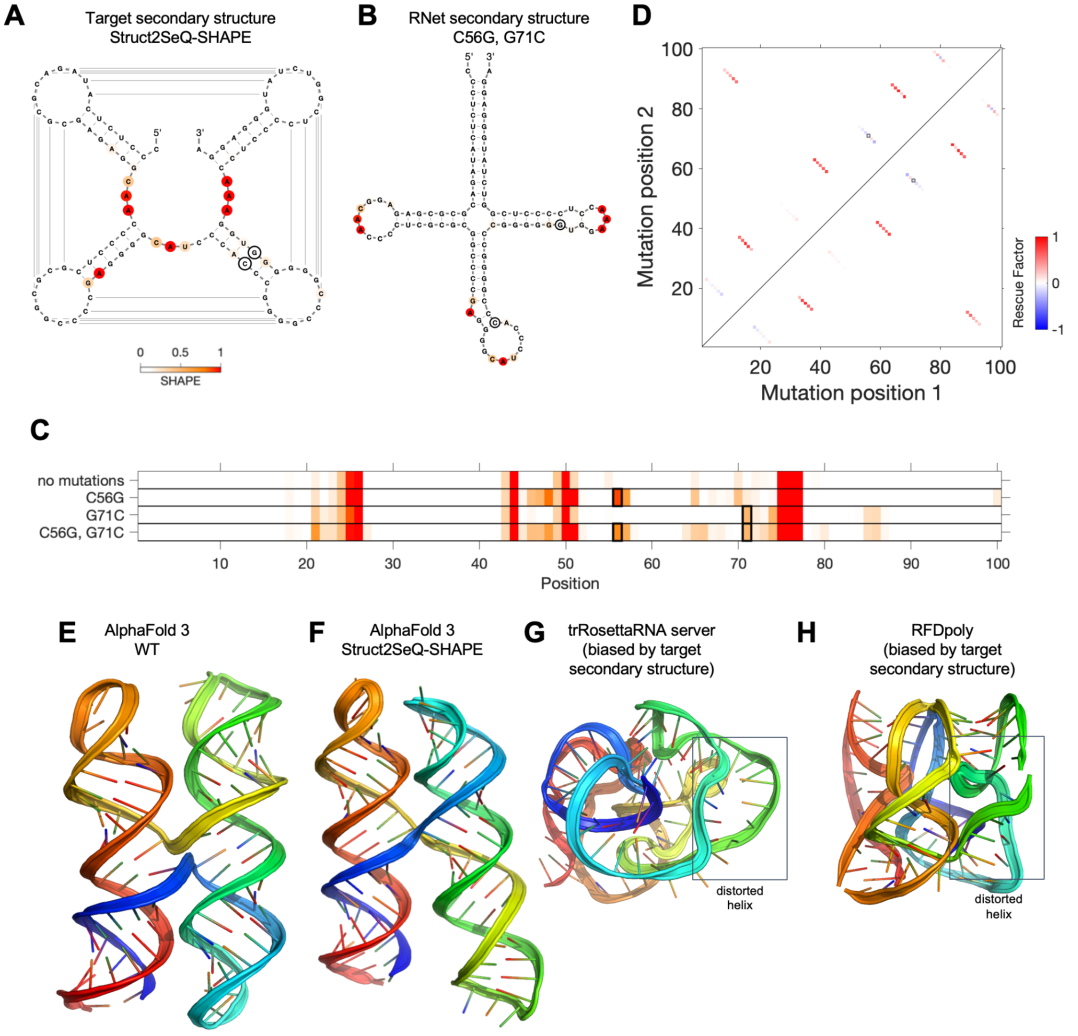

**Fig. S7. Incorrect base pairs in designs uncovered by mutate-map-rescue.** (**A**-**B**) SHAPE data for Struct2SeQ-SHAPE design with highest OpenKnot score for Round 3 target P16 (AK_PK100-3), colored onto (**A**) target pseudoknot secondary structure and (**B**) RNet-SS-predicted secondary structure. Circles highlight C56 and G71, which should form a base pair but are predicted to be unpaired by RNet-SS. (**C**) SHAPE profiles were perturbed at and outside the site of mutations C56G and G71C in single mutants, and these disruptions were not rescued in double mutant (bottom row). (**D**) Summary of rescue factors testing all target base pairs. Only RNet-SS -predicted pairs showed evidence for rescue upon compensatory mutagenesis (red), not 56-71 or the other target pairs (light red/blue). (**E**-**F**) AlphaFold 3 models of the starting sequence and the Struct2SeQ-SHAPE design produce alternative, non-pseudoknotted secondary structure. (**G**-**H**) 3D models based on (**G**) trRosettaRNA and (**H**) RFDpoly, both with the target secondary structure provided as input, produced distorted helices (boxes highlight examples). These results suggest that the target secondary structure may not be compatible with a stereochemically feasible three-dimensional arrangement of the target helices, potentially exacerbated by the symmetry and shorter helix lengths of the target secondary structure (*58*).

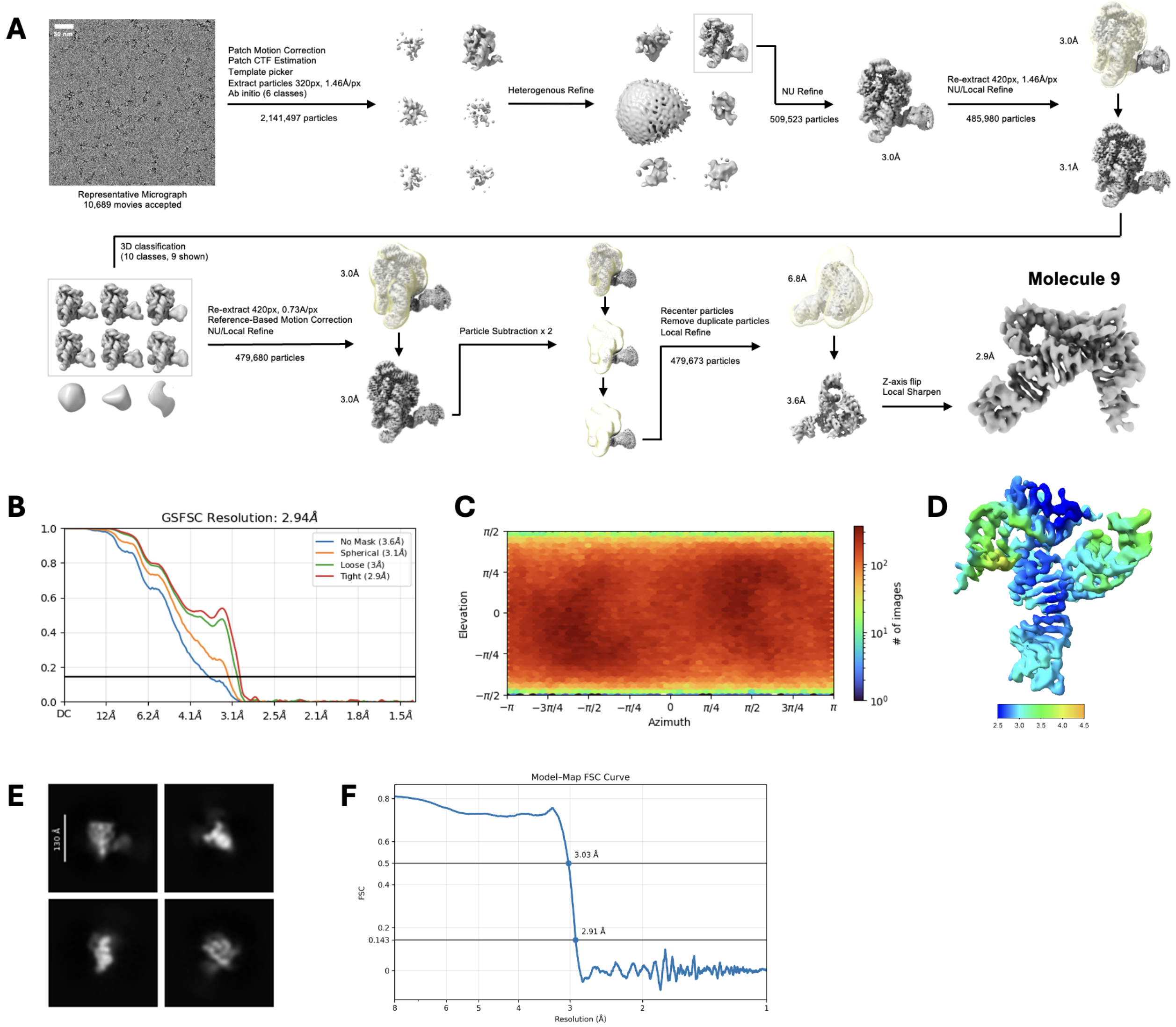

**Fig. S8. Cryo-EM data processing of gRNAde design ('Mol9'). (A)** Data processing workflow. **(B)** Gold Standard Fourier Shell Correlation curve. **(C)** Viewing direction distribution plot. **(D)** Local resolution heat map. **(E)** Representative 2D class averages of the scaffold attached to RNA insert **(F).** Model-map FSC curve.

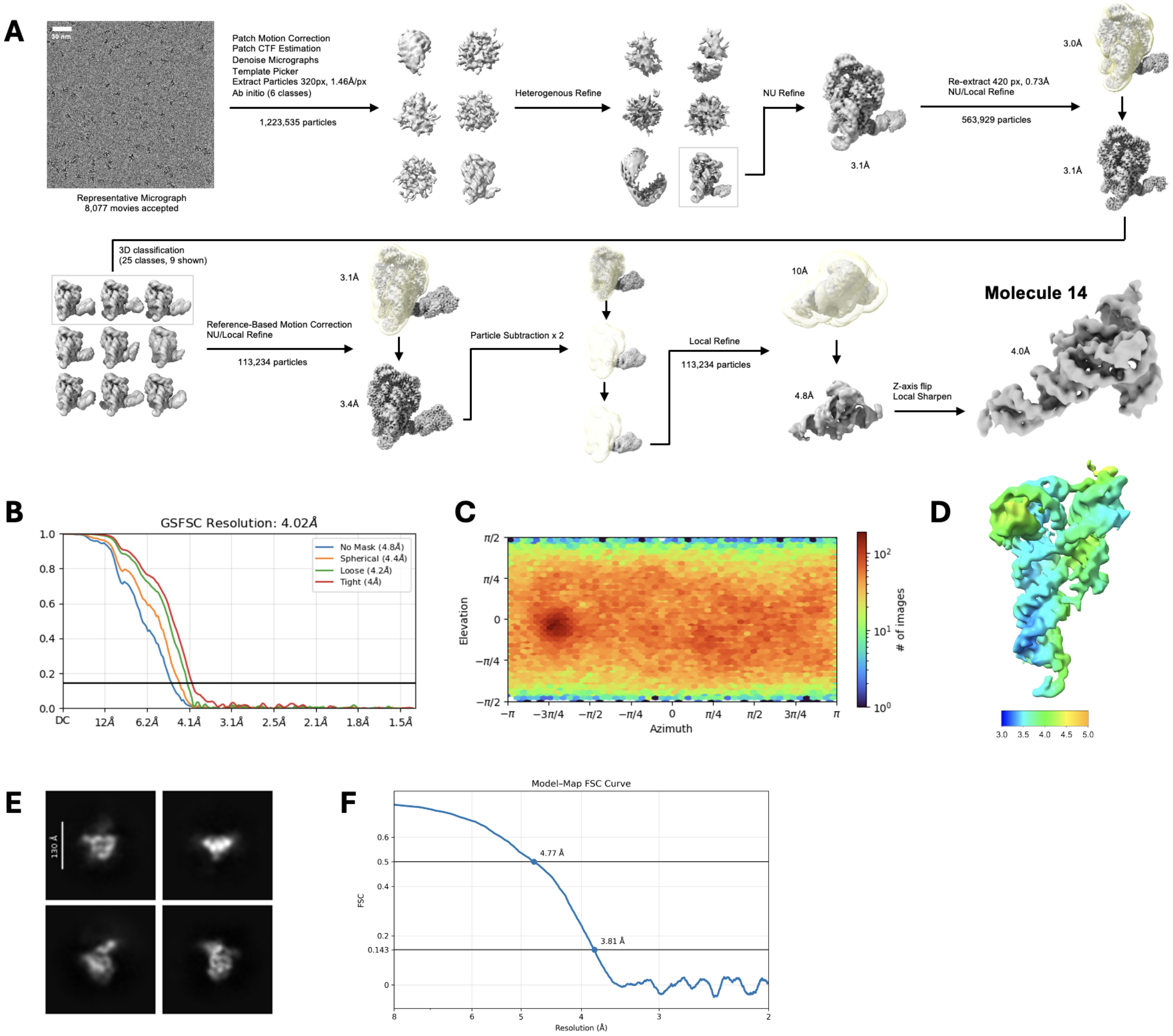

**Fig. S9. Cryo-EM data processing of MPNN-fixedbb design ('Mol14'). (A)** Data processing workflow. **(B)** Gold Standard Fourier Shell Correlation curve. **(C)** Viewing direction distribution plot. **(D)** Local resolution heat map. **(E)** Representative 2D class averages of the scaffold attached to RNA insert **(F).** Model-map FSC curve.

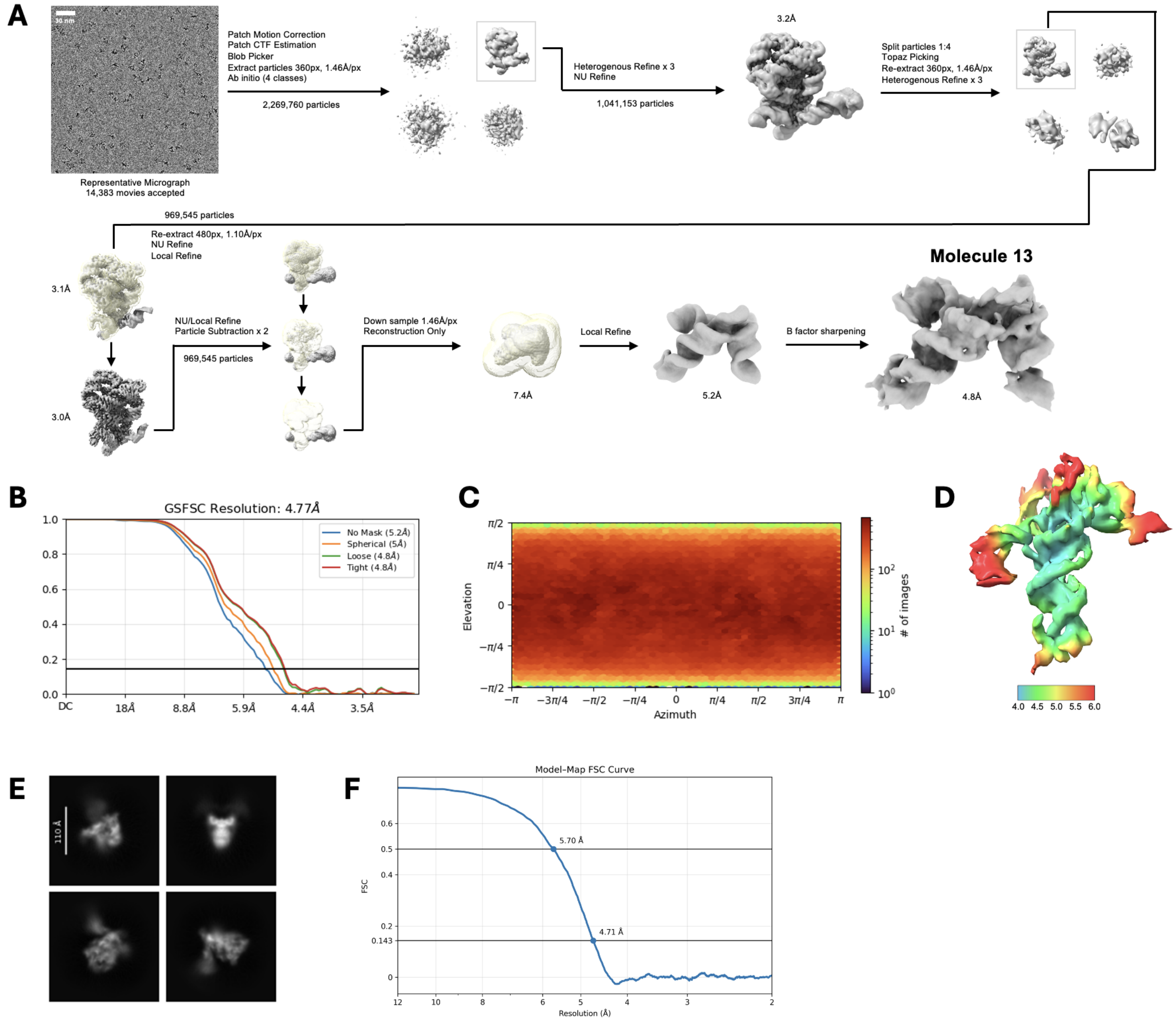

**Fig. S10. Cryo-EM data processing of Struct2SeQ-SHAPE design ('Mol13'). (A)** Data processing workflow. **(B)** Gold Standard Fourier Shell Correlation curve. **(C)** Viewing direction distribution plot. **(D)** Local resolution heat map. **(E)** Representative 2D class averages of the scaffold attached to RNA insert **(F).** Model-map FSC curve.

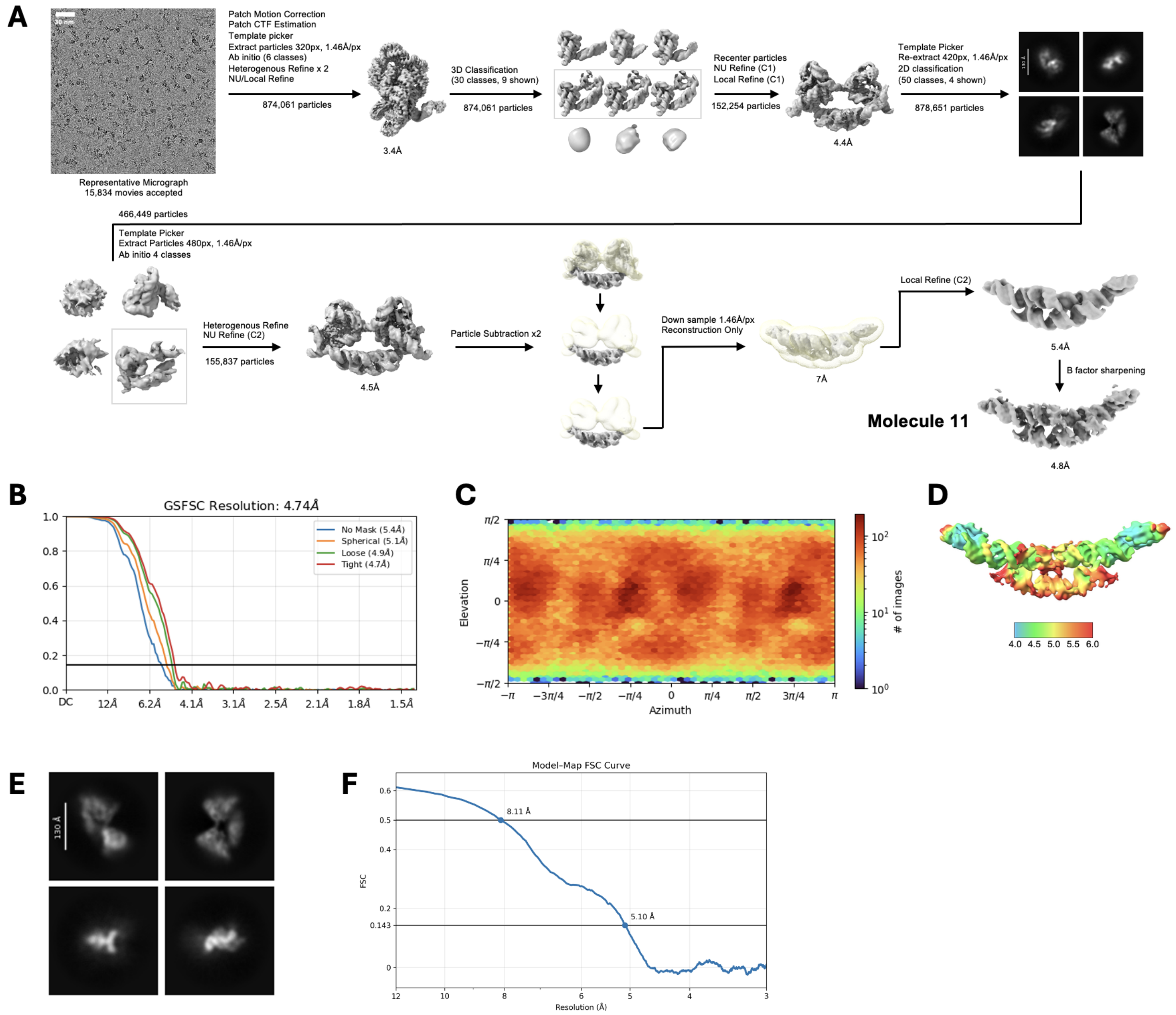

**Fig. S11. Cryo-EM data processing of MPNN-RFdiff design dimer ('Mol11'). (A)** Data processing workflow. **(B)** Gold Standard Fourier Shell Correlation curve. **(C)** Viewing direction distribution plot. **(D)** Local resolution heat map. **(E)** Representative 2D class averages of the scaffold attached to RNA insert **(F).** Model-map FSC curve.

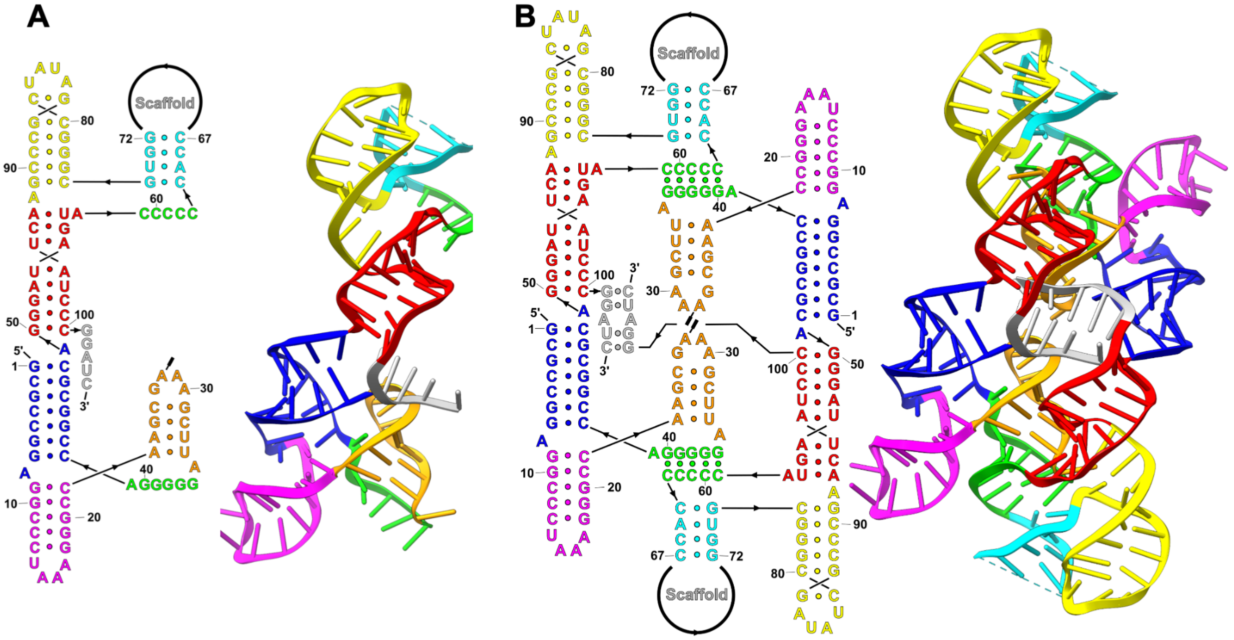

**Fig. S12. Secondary structure and Cryo-EM 3D structure of MPNN-RFdiff design dimer.** (**a**) Monomer extracted from symmetric dimer, (**b**) Complete dimer. Gray regions mark site of insertion of group II intron scaffold and extra nucleotides at 3´ end derived from BamHI restriction site.

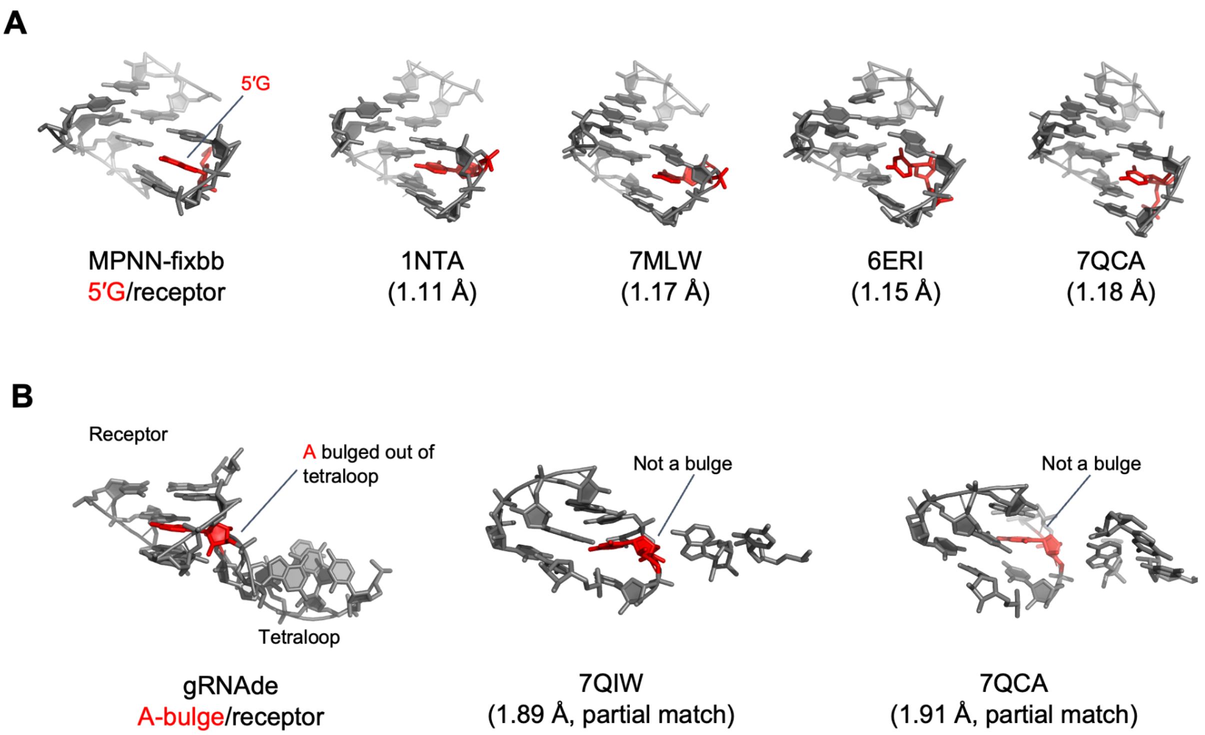

**Fig. S13. Best matches of AI design tertiary contacts to prior databases with ARTEM.** (**a**) Best ARTEM hits to 5´G/receptor of MPNN-fixbb design recover base pairing pattern as well as intercalation into receptor by nucleotide (red) from separate strand than receptor. (**b**) Best ARTEM hits to A-bulge/receptor of gRNAde design are partial matches with poor RMSD and do not have the intercalated nucleotide bulged out of a separate loop.

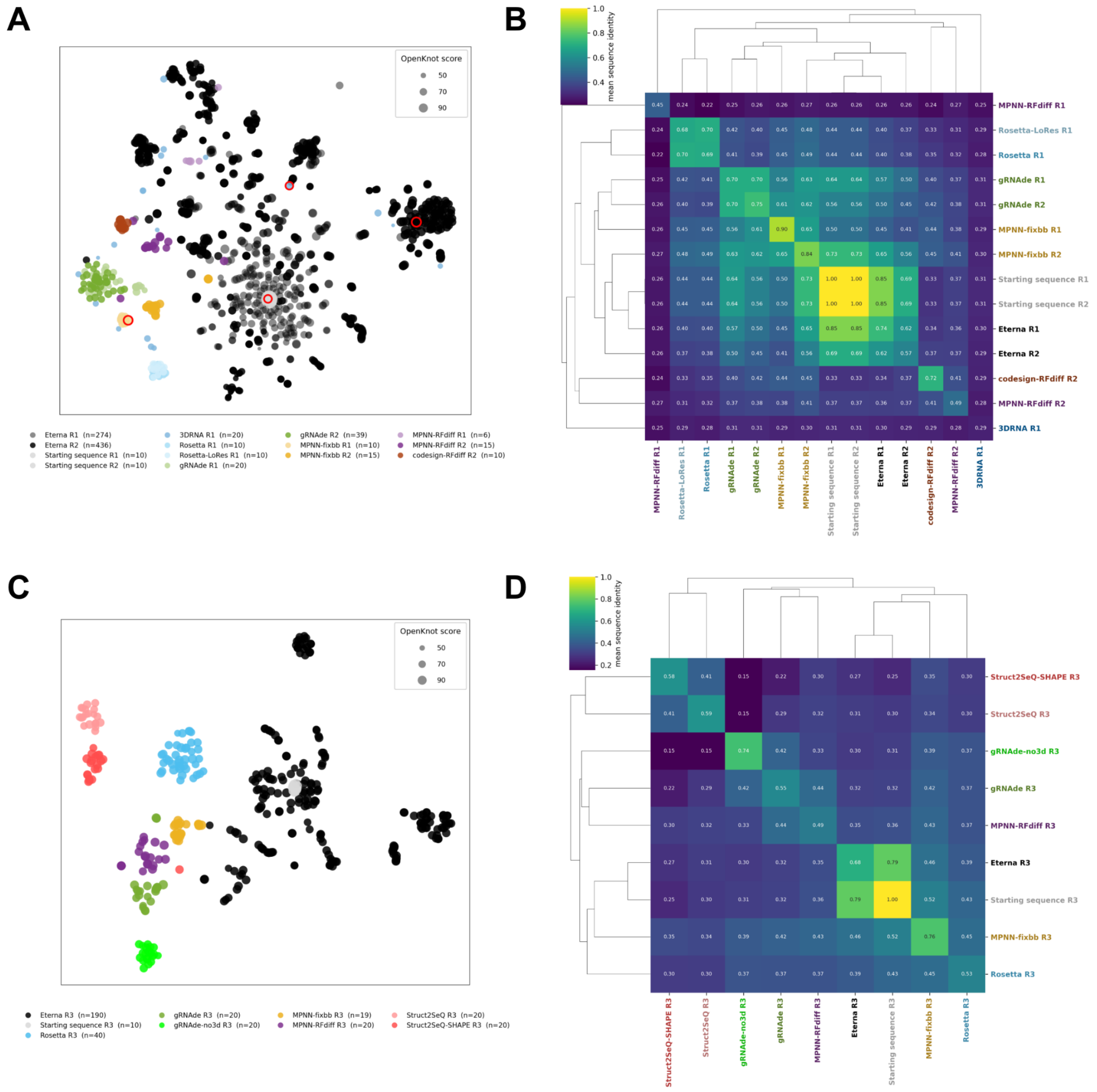

**Fig. S14. Sequence-space analysis of OpenKnot designs.** (**A**, **C**) t-SNE projections of all designs for targets W03 (Round 1, Round 2; $N = 885$) and P20 (Round 3; $N = 359$), respectively, colored by design method; marker area scales with OpenKnot score. Round 1 W03 designs are shown in lighter shades; Round 2 in full color. (**B**, **D**) Hierarchically clustered heatmaps of mean pairwise sequence identity within and between groups for W03 and P20, respectively (average linkage, optimal leaf ordering). Each AI method (colors) occupies a tight cluster in sequence space (within-method identities 0.70–0.90 for most methods), while Eterna designs (black) span a broader region anchored near the natural starting sequence. AI methods sample largely disjoint regions from one another (cross-method identities 0.40–0.50). Note that t-SNE preserves local cluster structure but distorts global distances; pairwise sequence identities in panels (**B**, **D**) are the quantitative measure of similarity.

1

| Player ID | Player Name | Real name | Designs |
|---|---|---|---|
| 8627 | Eli Fisker | Eli Fisker | 2461 |
| 64544 | AndrewKae | Andrew Kaechele | 2238 |
| 215715 | Astromon | William Dunlap IV | 2008 |
| 282879 | ucad | | 1877 |
| 48170 | Jieux | | 1872 |
| 273003 | mjt | Mary Joanne Travers | 1857 |
| 32627 | jandersonlee | Jeff Anderson-Lee | 1694 |
| 42833 | JR | | 1471 |
| 255372 | Father_Z | Zbigniew Grabara | 1412 |
| 292973 | jenni111 | Jenni Exley | 1165 |
| 265 | spvincent | Stephen Vincent | 1137 |
| 237588 | tone | | 851 |
| 223393 | dl2007 | | 716 |
| 32487 | stevetclark | Steven Clark | 478 |
| 246020 | DigitalEmbrace | Jill Townley | 467 |
| 272129 | Merida | Shannon Locke | 214 |
| 228503 | dizzywings | | 148 |
| 423707 | str1ngs | | 120 |
| 244068 | Mikiea | | 80 |
| 263010 | amybarish | Amy Barish | 79 |
| 337282 | Norbert13 | | 70 |
| 225146 | worseize | | 63 |
| 282312 | OSC | Osman Salih Ceyhan | 52 |
| 386401 | KagemiSky | | 50 |
| 280913 | voltor | Teresa Atienza | 30 |
| 196596 | whbob | Bob Conroy | 30 |
| 248916 | SuperD333 | | 28 |
| 388605 | ilikebball | Arghyo Sarkar | 27 |
| 241903 | Anamfija | Slava Fazio | 26 |
| 355382 | thumbs_up | Alexandra Sloman | 19 |
| 37055 | BugacMan | | 15 |
| 230687 | kriss888 | | 14 |
| 390683 | ThisNameIsNotAPun | | 14 |
| 328103 | d3admanu | | 13 |
| 26880 | Poll na gColm | | 12 |
| 245344 | DogeSka | | 12 |
| 260362 | TheInverted | Henry P. Dennis | 12 |
| 129706 | Trentis1 | | 11 |
| 349587 | arcpolaris | | 11 |
| 254029 | hi1000 | | 10 |

| | | | |
|---|---|---|---|
| 337520 | Creativa | | 8 |
| 342053 | AdityaL | | 8 |
| 338108 | KangLeTian | | 8 |
| 354798 | AliceDG | | 8 |
| 57675 | Omei | | 8 |
| 236173 | holoaaron | | 6 |
| 309589 | Juster | | 6 |
| 355481 | maestrocole | | 6 |
| 360357 | Blueberries-Are-Not-Blue | | 6 |
| 256114 | asiaa | | 5 |
| 379305 | StargazerCGM13 | | 4 |
| 359696 | man_the_stan | Stan Soo | 4 |
| 330968 | AkiraHatsuko | | 3 |
| 148353 | aet36 | Andy Romano | 3 |
| 348141 | Flamin'Homosapien | | 3 |
| 299221 | CoopPlayerLP | | 3 |
| 359113 | eleseduncan | | 3 |
| 231740 | David Dockrell | | 3 |
| 382338 | leonardcarter71810 | | 3 |
| 243727 | gootboy54 | Josh Lavelle | 3 |
| 368380 | terraofearth | | 3 |
| 268069 | Wipf0 | | 2 |
| 343777 | JoltCode | | 2 |
| 340277 | tokiWren | | 2 |
| 11775 | starryjess | | 2 |
| 324670 | henry552 | | 2 |
| 294279 | ElMounRo | | 2 |
| 348556 | ALP | | 2 |
| 356012 | BanBan | | 2 |
| 355853 | Anushka_Verma | | 2 |
| 382415 | s127679 | | 2 |
| 352426 | EliteZeldaGamer | | 2 |
| 412002 | Sealman | | 2 |
| 398783 | KARATENAT | | 2 |
| 342806 | Giahuy2004 | | 1 |
| 282833 | Tbuhlig | | 1 |
| 318126 | anitaavg | | 1 |
| 340151 | LightningHolt | | 1 |
| 265289 | Euclides | | 1 |
| 340899 | mr33h | | 1 |
| 272147 | XGO | | 1 |
| 341473 | Thomas_C | | 1 |

| | | | |
|---|---|---|---|
| 136909 | wookieetank | | 1 |
| 320985 | sgeldhof | | 1 |
| 376808 | bruhnugget | | 1 |
| 356677 | nkvamaks | | 1 |
| 360918 | Ronaldo/GOAT | | 1 |
| 274312 | FRAG3R | | 1 |
| 375485 | unetoilebrille | | 1 |
| 295555 | brainbow | | 1 |
| 359109 | Yak | | 1 |
| 379138 | WingedDungBeetle | | 1 |
| 363214 | Nico_F | | 1 |
| 290933 | febos | Eugene F. Baulin | 1 |
| 415540 | Hannah~ | | 1 |
| 41515 | Rivalium | Rebecca L. Taccone | 1 |
| 387919 | Luka3231 | | 1 |

**Table S1. Eterna Participants.** Eterna players who contributed designs for the four rounds of the OpenKnot AI challenges; all are independent scientists affiliated with the Eterna Massive Open Laboratory, USA. Major contributor Jill Townley can be contacted on behalf of the group.

1

| Method | Eterna | 3DRNA | gRNAde, gRNAde-no3d | MPNN-fixbb, MPNN-RFdiff, codesign-RFdiff | Rosetta, Rosetta-LoRes | Struct2SeQ, Struct2SeQ-SHAPE |
|---|---|---|---|---|---|---|
| **Tasks with notable performance** | Top method, Round 1<br>Top method, Rounds 2-4* | (discontinued after poor performance in Round 1) | Top AI method (Z-score), Round 3*<br>Highest resolution cryo-EM | Top AI methods, Round 2* (top overall by Z-score*)<br>Top M2R score, Round 3*<br>Cryo-EM | (Pre-deep learning baseline) | Top AI method in Round 3*<br>Top AI method in Round 4**<br>Cryo-EM |
| **Model architecture** | Not automated (on-line community of human participants) | 3D convolutional neural network | Graph neural network, autoregressive RNA language model | MPNN: Graph neural network<br>RFdiff (renamed RFDpoly): Diffusion generative model | Simulated annealing guided by hand-crafted energy function | Deep-Q reinforcement learning |
| **Secondary structure (2D) input** | Requires 2D input | No | Optional (Initial gRNAde variant in Rounds 1 and 2 required 3D backbone input) | Optional 2D input to RFdiff | No | Requires 2D input |
| **3D Structure input** | Optional; in-game visualization of single model backbone | Requires 3D backbone input | Optional (Initial gRNAde variant in Rounds 1 and 2 required 3D backbone input) | MPNN-fixbb requires backbone input | Requires 3D backbone input | No |
| **Training data** | Human learning on prior Eterna challenges, including free-form pseudoknot design. | Prior PDB | Prior PDB (October 2023 cutoff) | Prior PDB | None (not machine learning) | "Playouts" on target secondary structures, guided by RNet |
| **Filtering** | Not automated | None | RNet SHAPE predictions; RNet-SS 2D structure (gRNAde perplexity, EternaFold also used in Rounds 1 and 2) | RNet SHAPE predictions; RNet-SS 2D structure (EternaFold used in Round 1) | None | RNet-SS 2D structure; RNet SHAPE predictions (for Struct2Seq-SHAPE) |
| **Extensibility** | Currently limited to RNA 2D structures | Currently limited to RNA 3D fixed backbone | Multi-structure RNA ensembles, Supports 2D-only or 3D backbone + 2D conditioned design. | DNA, nucleic-acid/protein complexes | Currently limited to RNA 3D fixed backbone | General objective functions (requires re-training) |

* Data did not discriminate 'top' method from next best methods with statistical significance, after correction for multiple hypothesis testing.
** Struct2SeQ and Struct2SeQ-SHAPE designs for Round 4 were submitted after release of experimental results for other design methods in Round 4.

**Table S2.** Summary of design methods in OpenKnot AI challenge

| OpenKnot AI round number | Eterna OpenKnot internal round number | Number of Targets | Maximum length (nt) | Eterna link | Number of Eterna participant designs | Number of AI designs |
|---|---|---|---|---|---|---|
| 1 | OpenKnot Round 4 | 17 | 100 | https://eternagame.org/labs/13189441<br>https://eternagame.org/labs/13196868<br>https://eternagame.org/labs/13200403<br>https://eternagame.org/labs/13212579<br>https://eternagame.org/labs/13214425<br>https://eternagame.org/labs/13195459<br>https://eternagame.org/labs/13227720<br>https://eternagame.org/labs/13235921<br>https://eternagame.org/labs/13247192<br>https://eternagame.org/labs/13251661<br>https://eternagame.org/labs/13256720<br>https://eternagame.org/labs/13274774<br>https://eternagame.org/labs/13278128<br>https://eternagame.org/labs/13206733<br>https://eternagame.org/labs/13292843<br>https://eternagame.org/labs/13298184<br>https://eternagame.org/labs/13307077 | 5790 | 1162 |
| 2 | OpenKnot Round 6 | 17 | 100 | https://eternagame.org/labs/13389097 | 7292 | 1086 |
| 3 | OpenKnot Round 7a* | 20 | 100 | https://eternagame.org/labs/13612323 | 3966 | 3205 |
| 4 | OpenKnot Round 7b* | 20 | 240 | https://eternagame.org/labs/13612324 | 3396 | 2620 |

* selected in "OpenKnot Sprint: Players pick the Targets": https://eternagame.org/labs/13618132

**Table S3.** Summary statistics for four rounds of the OpenKnot AI challenges

| | | | | | | Round 1 | | Round 2 | |
|---|---|---|---|---|---|---|---|---|---|
| **ID** | **Title** | **Length** | **Dot-bracket** | **Starting sequence** | **PDB source** | **Number of Eterna participant designs** | **Number of AI designs** | **Number of Eterna participant designs** | **Number of AI designs** |
| W01 | 6XRZ SARS-CoV-2 FSE | 88 | (.((......))..).((((((((((.(([[[[[)))))))))))))((((((((..(....).))).)))))).....]]]]].... | GUUUUUAAACGGGUUUGCGGUGUAAGUGCAGCCCGUCUUACACCGUGCGGCACAGGCACUAGUACUGAUGUCGUAUACAGGGCUUUUG | RCSB PDB | 324 | 69 | 432 | 79 |
| W02 | 7KD1 THF riboswitch | 89 | ((((....((((((((((((......(((((((...[[[[[...))))))))).....((....))))).))))))))).]]]]].)))). | GGAGAGUAGAUGAUUCGCGUUAAGUGUGUGUGAAUGGGAUGUCGUCACACAACGAAGCGAGAGCGCGGUGAAUCAUUGCAUCCGCUCCA | RCSB PDB | 303 | 75 | 435 | 78 |
| W03 | 3Q3Z c-di-GMP-II switch | 74 | (((((.(((......(((((((((..(..{[[[[[[[)))))))))))...)))...]]]]]}.]]..))))).. | CGCGGAAACAAUGAUGAAUGGGUUUAAAUUGGGCACUUGACUCAUUUUGAGUUAGUAGUGCAACCGACCGUGCU | RCSB PDB | 275 | 76 | 436 | 79 |
| W04 | 3T4B HCV IRES | 84 | (((((((((((.((((((.[((((....)))))]))))))[))((((..].))))((..[[[[[.)))))))))))).]]]]]... | CCUCCCGGGAGAGCCGCUAAGGGGGAAACUCUAUGCGGUACUGCCUGAUAGGGUGCUUGCGAGUGCCCCGGGAGGUCUCGUAGA | RCSB PDB | 295 | 75 | 425 | 79 |
| W05 | 2AE KL | 90 | .(((((((((((((((((((..[[[[[[.)))))(((....)))(((....))))))))))))))))((((((..]]]]]].))))))). | CCUGGGAACCUCCUGUGCUGAAGCGCGCACGGCAACCGAAAGGUGCGGAAACGCUGGGGGGUUCCUGGAUGCUGAAGCGCGCACGGCAUU | Cody Geary | 360 | 76 | 443 | 75 |
| W06 | 4RZD preQ1 class 3 | 100 | ((((......[[[[[[...))))(((((((((........)))))))))....(((((((((((........))))))).))).).]]]]]].......... | GAGCAACUUAGGAUUUUAGGCUCCCCGGCGUGUCUCGAACCAUGCCGGGCCAAACCCAUAGGGCUGGCGGUCCCUGUGCGGUCAAAAUUCAUCCGCCGGA | NA | 335 | 57 | 458 | 74 |
| W07 | 4PLX Malat1 | 76 | .[[[[[..........((((((((....)))))))((((((((((((..]]]]]...........))))))))))) | GGAAGGUUUUUCUUUUCCUGAGGCGAAAGUCUCAGGUUUUGCUUUUUGGCCUUUCUUAAAAAAAAAAAAAGCAAAA | NA | 351 | 60 | 447 | 38 |

| | | | | | | | | | |
|---|---|---|---|---|---|---|---|---|---|
| W08 | 2M8K telomerase | 48 | [[[[[......(((((((((<br>(((]]]]].....).).)))<br>))))))). | GGUUUCUUUUUAGUGAUUU<br>UUCCAAACCCCUUUGUGCA<br>AAAAUCAUUA | RCSB PDB | 317 | 81 | 488 | 78 |
| W09 * | Eterna ZG 113 | 201 | .(((((.......((((((((<br>(............{{{{{{<br>{.......)))))))))....<br>.((((..}}}}}}}.))<br>)))))))..............<br>.(((((.(((...(((((((<br>((............[[[[[[..<br>......))))))))).......<br>.....]]]]]].))).))))<br>)............... | ACUGCGACAGAAAAGAAGC<br>GUGAAACAAAAUGAAUCUC<br>GGCAAAGCUACGCGCUUCU<br>ACAUACCAGCAGCCGAGAA<br>CUGGCGCAGAUUACAAACG<br>CAGAC | NA | 313 | 59 | 424 | 37 |
| W10 | Eterna UC2199 | 100 | (((((((..............(<br>((((......{{{{{{..<br>...)))))......((((((<br>.......}}}}}}.....)<br>)))))....)))))))).... | GAGCCCGUUGCAUUUUAUU<br>UUCGAGCUUUCUUGCCGCC<br>UUCUUGCUCGUAUUGCUGG<br>ACCGUUUCUUUGGCGGCUU<br>CUUCGGUCCUAUUCGGGCU<br>CGAGA | NA | 376 | 64 | 431 | 40 |
| W11 | Eterna AK-PK_380 | 100 | ((.(({{{{{{{{.....<br>((((((.......)))).)))<br>).))......((((((((((<br>((((((((...}}}}}}<br>}))))))))...)))))))<br>)).... | GCUGGCCUGCGCACGAUCU<br>GACUGCCGUGAAGUCUAGC<br>CUGCGUCAGAACUAGUCUC<br>CAUCAAUCAAAGCGCAGGG<br>AUUGAUGUACGAGACUAGU<br>ACACA | NA | 307 | 56 | 400 | 40 |
| W12 | Eterna S. aureus | 100 | ...(((((.{{{{{{.)<br>))))....(((((((......<br>.}}}}}}...........(<br>((.((((........)))).)<br>))((....))....)))))))<br>. | CUACCUUUCCAUGCUAAAA<br>GGUUCAAUGAUUGGUUAGA<br>AAGCAUGUCAAAUAAUGUC<br>GUUCGAUCAAAAGGUAUCG<br>UAUGGUUAGCACAAUACAA<br>UCAUG | NA | 376 | 60 | 408 | 40 |
| W13 | Eterna S. tuberosum | 100 | .((((((((((.((((((..<br>{{{{{.....(((....))<br>)............))))))....<br>.......(((.......)))}<br>}}}}})))))))..))). | UGGCCUACCAUACUAUCCG<br>UUAUAUAAAUAGGCAAAUG<br>CUCUUUUCCAUUAUGGAUA<br>GCAAUAGUAUGACCGAUCA<br>UUGUGGGUAUAAUGGUAGA<br>UGCCC | NA | 363 | 60 | 419 | 40 |
| W14 | 4OQU SAM I/IV switch | 97 | (((((....((((....))))<br>(((..(((((((.[[[[[.))<br>).))).)))..(((((....) | GGAUCACGAGGGGGAGACC<br>CCGGCAACCUGGGACGGAC<br>ACCCAAGGUGCUCACACCG<br>GAGACGGUGGAUCCGGCCC | RCSB PDB | 344 | 75 | 394 | 77 |

| | | | | | | | | | |
|---|---|---|---|---|---|---|---|---|---|
| | | | ))))))))).((((....))<br>))........]]]]]. | GAGAGGGCAACGAAGGUCC<br>GA | | | | | |
| W15 | 7KGA donggang dumbbell | 89 | ((((((((((((((((((<br>(((...[[[[.))))))).)<br>.)))).))))).((.......<br>...))......)))))........<br>..]]]] | GGGGGCCAGAUGUCAUGUC<br>UCUCAAGCCUAGGAGACAC<br>UAGACACUCUGGACUAUCG<br>GUUAGAGGAAACCCCCCCA<br>AAAAUGUAUAGGC | RCSB PDB | 331 | 66 | 394 | 76 |
| W16 | 1DRZ HDV ribozyme | 72 | (((((((...[[[[[[(((.<br>[[.....))))))))))]].<br>...((((..........)))).<br>....]]]]]] | GGCCGGCAUGGUCCCAGCC<br>UCCUCGCUGGCGCCGGCUG<br>GGCAACACCAUUGCACUCC<br>GGUGGCGAAUGGGAC | RCSB PDB | 398 | 77 | 391 | 79 |
| W17 | 7QR4 CPEB3 | 69 | [[[[[[...((((((((((.<br>......))).]]]]]]..(((<br>((..........)))))....)<br>)))))) | GGGGGCCACAGCAGAAGCG<br>UUCACGUCGCAGCCCCUGU<br>CAGCCAUUGCACUCCGGCU<br>GCGAAUUCUGCU | RCSB PDB | 422 | 76 | 467 | 77 |

*The target structure for the round 1 submission changed after the W09 Target opened. Both structures were used for computing OpenKnot score (maximum score assigned to designs).

**Table S4.** Targets for OpenKnot AI Rounds 1 and 2

1

| ID | Title | Source | Length | Dot-bracket | Sequence | PDB source | Number of Eterna participant designs | Number of AI designs |
|---|---|---|---|---|---|---|---|---|
| P01 | Thermotoga petrophila fluoride riboswitch | 4ENA | 52 | .[[[[{.((((((]]]]......(((((....)))))}.[.))))))].... | GGGCGAUGAGGCCCGCCCAAACUGCCCUGAAAAGGGCUGAUGGCCUCUACUG | trRosetta | 221 | 160 |
| P02 | ZTP switch | 5BTP | 75 | .....(((((((..(.[[[[[...((((......))))..)..)))))))........(((..]]]]]..))). | UAUCAGUUAUAUGACUGACGGAACGUGGAAUUAACCACAUGAAGUAUAACGAUGACAAUGCCGACCGUCUGGGCG | trRosetta | 224 | 160 |
| P03 | pfl switch | 4ZNP | 73 | .((((((((..(.[[[[[....((((....))))..)..))))))))........(((..]]]]]..)))... | GGGAUACAGGACUGGCGGAUUAGUGGGAAACCACGUGGACUGUAUCCGAAAAAAAGCCGACCGCCUGGGCAUC | RNASolo | 220 | 160 |
| P04 | sgRNA | 5X2G | 93 | ....................((((((((((((....))))))))))))..((((...[[[[..))))..((((.......)))).....]]]] | GGAAAUUAGGUGCGCUUGGCGUUUUAGUCCCUGAAAAGGGACUAAAAUAAAGAGUUUGCGGGACUCUGCGGGGUUACAAUCCCCUAAAACCGC | RCSB PDB | 237 | 160 |
| P05 | PreQ1-II switch | 2MIY | 59 | ((((((((.....[[[[[[.)))))))).........((((....))))..]]]]]].. | GCUUGGUGCUUAGCUUCUUUCACCAAGCAUAUUACACGCGGAUAACCGCCAAAGGAGAA | RCSB PDB | 232 | 160 |
| P06* | Grapevine leafroll-associated virus - 2 | PKB271 | 75 | .((((.....((((.......))))[[[[[[))))((((((....))))))..................]]]]]]<br>.((((.....((((.......))))[[[[[.))))((((((....))))))(((((((.....)))))))]]]]] | ACGCCAAAAUCCAAUUAAAGUUUGGGACCUAGGCGGGCCUCUUACGAGGCUAACUUAUCGACAAUAAGUUAGGUC | trRosetta | 204 | 160 |
| P07 | Turdivirus 3 | PKB366 | 80 | ..(((((..[[[[[)))))..............((((((.((((((((....)))))))).))))))).....]]]]].......... | UCCGUCCAAGCCGCGGACGUUAAACUGUGGCACGGUUGUGCCUAGGUGCAACCCUGCUACUAAUGGCGGUACCCCUGCCC | trRosetta | 206 | 160 |
| P08 | Poa semilatent virus | PKB164 | 96 | .((((((((..((((((..[[[[[.))))))....((((((......))))))))))))))).((((((.....))))))........]]]]].... | AUUGGUAUGUAAGCUACUUCUUCCAGUAGCUGCGUCAUAACAUCAAGGUUAUGCAUACUGAGCCGAAGCUCAGCUUCGGUCCUCCAAGGAAGACCA | trRosetta | 212 | 160 |

| | | | | | | | | |
|---|---|---|---|---|---|---|---|---|
| P09 | E. coli | PKB129 | 68 | ..((....[[[[[[..((((((((((((.....)))))))))))))...[[..))]].....]]]]]]. | GCGUAAAUGUCGACUUGGAGGUUGUGCCCUUGAGGCGUGGCUUCCGGAGCUAACGCGUUAAGUCGACC | FARFAR2 | 202 | 160 |
| P10 | Diplonema papillatum | PKB341 | 76 | (((((((......[[[.[[[[[(((.[[....))))))))))]]...((((((.....)))))......]]]]]]] | GAGGGACAAGAAUCUGACCUGCACCUCCUCGUGGUGUCCCUCGGAAACGUGCUCAACGCGCGGCCGACGCAGGCAG | FARFAR2 | 196 | 160 |
| P11 | PN.v282 | EcceruElme | 100 | ((((((((((.(((......))).)))))))[[[[[[((((((((..(((((((......))))...))).))))))))...))))).......]]]]]] | CUGCCGACGCCCUGUGUUGAGGCGUGGUGUCGCCUCGAGUCAGUCGUUGAGAUCCGAAGAAGGGUGUACUUCUGACUGAUCGAGCAGAAGUGAGUCGAGG | trRosetta | 183 | 165 |
| P12 | mod of f67 | hi1000 | 100 | ......((((((.((((((....))))))(((((....[[[[[[[[[(((((((....)))))))).....)))))..))))))..]]]]]]]]]...... | GAAAAGAGCGUAAUCGCGUCUUGACGUGAGUGAGGCUAGACUGUAGACUAUCCCAAGUGGGAUAGCAUAACUCACAAUACGCUGAUCUACAGUCAAAAGA | trRosetta | 186 | 160 |
| P13 | UC2414 | ucad | 100 | (((((((..[[[[[))))))))..((..((....))..))..(((((.........)))))...(((((((...........]]]]]]..)))))))... | GGAAGACAACUCGCGGUCUUCCAAGUACUCGAAGGAAGGCGAGACGCACGAGUGCAGCGUCAAAUGACACCACGUCAUCCAACGCGAGAAGGUGUCAGAA | FARFAR2 | 184 | 160 |
| P14 | SV_j116 | spvincent | 100 | .((((((((((([[[[[[[[(((((....)))))........))))))))))......(((((((((((((..]]]]]]]]..))))))))))))))....... | GAGGUGAUAAUCUGAUAGGAGGCUGAGAAGUCUGAAGGUAAAUUAUCGUCUAAUUAAAUCUGGUUUAAGUACCUAUCAGAUCUUAGACUAGAUGAAGAUA | trRosetta | 189 | 160 |
| P15 | UC2458 | ucad | 100 | (((((.[[[[[[[[))))))...(((((((..{{{{{{))))))))..]]]]]]]].......((((((((((((........)))))))))))..}}}}}}. | GUGGCACGUGUGCAGCCACGAAGGUUGGUAACACCGGAUCAACCGAUGUACACGACGUCUCGGAAUGCCAUGUUGAGACGUAUGGUAUUCCAACCGGUGA | trRosetta | 183 | 160 |
| P16 | AK_PK100-3 | AndrewKae | 100 | .((((((([[[[[[[[[[)))))))...((((((]]]]][[[[[))))))).....((((((]]]]][[[[[))))))...((((((]]]]]]]]]]))))))). | AGUGUGCCCCGGACCUCGCACACAAAGGUUGGGAGGUGGGGACCAACCAAAAGCAUCGUCCCCGGCACCGAUGCAUAGCCUGGGUGCCCCGGGCCAGGCA | trRosetta | 173 | 160 |
| P17 | ZG21 | Father Z | 100 | (((......((((..(((((((((.....)))((((((...[[[[.))))))(((......]]]][[[[[[[..) | GGCCAGACAGCGUAACCGUGGCCCAAAAGGGGCGACGAGUACGGUAUCGUCGCGGAACUAUACCGGAUGGU | FARFAR2 | 168 | 160 |

| | | | | | | | | |
|---|---|---|---|---|---|---|---|---|
| | | | | ))))))))))))..]]]]]]].)))<br> | GAACCGCCACGGACGCAUCACCAUUUGCC | | | |
| P18 | Terminal 3 | DigitalEmbrace | 100 | .((((((((...((((((((((..[[[[[[.)))))))))))..(((((((.{{{{{{.))))))).))))))))))).....]]]]]].}}}}}}... | AGGUAAGUUGGAAGGUGUGAGGUGUUUCAUCAUCAUCUCACACCAAAGUACUUGCGAGAAAAAGUACUAACAACUUGCCAGCAAUGAUGAUUUCUCGACU | trRosetta | 187 | 160 |
| P19 | SV_r7_100_a7 | spvincent | 100 | (((((((((((((((((.....[[[[[[[[[[.[[[[[[[[[))))))))))]]]]]]]]].[[[[[[[.)))))))))]]]]]]]].]]]]]]]]]]. | GCUUCCGAAUGAUGACUCAAAUAGGUAAGUUGGAAGGUGUGAGGAGUCAUCAUCUCACACCUAAGUACUUGUUCGGAGGCCAAGUACUACCAACUUGCCA | FARFAR2 | 169 | 160 |
| P20 | Kissing multiloops | Eli Fisker | 100 | (((((((.(((((....)))))((((....)))).[[[[[.)))))))).(((((((.]]]]](((((....))))((((((....)))))).)))))))) | GGACGACAGUCUGCUUGCAGACCUCCGAAAGGAGAUGGCCAGUCGUCCUGGGUCAUCAGGCCAGUCGUUCGCGACGGUAGCGUGAGCUACCAGAUGACCU | FARFAR2 | 190 | 160 |
| Q01 | Guide RNA | 7YOJ | 174 | .((((.(...(.(((((..(.(.((....((((...(((.....)))...))))......)).).)..((((....)))).....)))))...)..(((.[[[[..)))).)))).(((((((((..((.....))..)))))))))...]]]].................... | GUCUGCCGAAGACGCCGCACGGAGCCUGGGCCGGAAUCGUAGAUCGAACGCGGCAUCGAAGCCCUGCAGCCCUUCGGGGCCAAGGCGGCGCAGCAAGCCUCUUUCAGGCGGCAGAGUCCUUUAGAGUGUGAGAGACACUCUAAAGGAAUGAAAGAGGGCGACACCCUGGUGAAC | RNASolo | 196 | 130 |
| Q02 | Nanobracelet | 7JRT | 134 | (((((((((((.......((((((.....((((((..[[[[[[.))))))))))))))))))))))))((((((((((.]]]]]].((((((.....((((((.........))))))))))))).)))))))))) | GGAUAUUCGACGGAGGCACCCAGGAACUACCGUUGAAGCUCGCACGACGGCCUGGGGUCGAGUAUCCCGGUAUUUGUCGCGAGCACUGAGGAACUACUGCUGAAGCCUCCACGGCAGCCUCAGGACAAGUACCG | RNASolo | 208 | 130 |
| Q03 | GLMS ribozyme | 3G9C | 140 | (((((..........)))))......[[[...((((((..]]]..............))))))((((((.[[[[[))))))...(((((((((.....((((((((....)))))))......)))))))))).]]]]] | GCACCAUUGCACUCCGGUGCCAGUUGACGAGGUGGGGUUUAUCGAGAUUUCGGCGGAUGACUCCCGGUUGUUCAUCACAACCGCAAGCUUUUACUUAAAUCAUUAAGGUGACUUAGUGGACAAAGGUGAAAGUGUGAUGA | RNASolo | 234 | 130 |
| Q04 | T-box | 6UFG | 166 | (((......((((......))))...)))((((((......((((.....)))).........)))))(((...[[[))). | GGCAUCGAUCCGGCGAUCACCGGGGAGCCUUCGGAAGAACGGCCGGUUAGGCCCAGUAGAACCGAACGGGUU | FARFAR2 | 183 | 130 |

| | | | | | | | | |
|---|---|---|---|---|---|---|---|---|
| | | | | ....]]].........{..(((((((....)))))))..((((((......}(((((((.(((((....)))))))))))))))))) | GGCCCGUCACAGCCUCAAGUCGAGCGGCCGCGCGAAAGCGUGGCAAGCGGGGUGGCACCGCGGCGUUCGCGCGAAAGCGUGGCGUCGUCCCCGC | | | |
| Q05 | Ligase ribozyme | 3HHN | 137 | ((((((((...[[[[[[.))))))))..............[[[[[(...).(.((((((((((((((.........)))))))..((((.]]]]])))))((.((((......)))).)))))))))).]]]]]]. | UCCAGUAGGAACACUAUACUACUGGAUAAUCAAAGACAAAUCUGCCCGAAGGGCUUGAGAACAUACCCAUUGCACUCCGGGUAUGCAGAGGUGGCAGCCUCCGGUGGGUUAAAACCCAACGUUCUCAACAAUAGUGA | FARFAR2 | 185 | 130 |
| Q06 | Taura IRES | 5JUP | 201 | ...(((((((..............((((((........[[[[[..........))))).........)))))))[[[[.(...........)..(((((]]]](.((......)).)..)))))......]]]]]..(((((.(((((((.[[[[[)))).))).((((((....))))))))))........]]]]]...... | AAACUCCAUGUAUUGGUUACCCAUCUGCAUCGAAAACUCUCCGAACACUAGGUGCAGUAAGGCUUUCAUGGAGUGGUUUGCUAUUUAGCGUACGUGUACCAUAGGCAGCCCCAAAAACACGUGUGAGGAGAAAGUCCCAGUCACUUUGGGCAAAGUAGACAGCCGCGCUUGCGUGGUGGGACUUAAUUAAUGCCUGCUAAC | trRosetta | 160 | 130 |
| Q07 | Broad bean mottle virus | PKB135 | 117 | ..(((((((..((((((..[[[[[[))))))......(((((.((((((.....))))))....))))))))))))(((((((((....)))))))))..........]]]]]].... | AAGAUAUCUGUACCGUCUUCUCCCCGACGGUUUCAAUCCAUAGCUUCACGUUGUGUAGCGUAGGUGGAAGGUAUCGUGUGGUGUAUAAACACCACAUAGUCUCUUAGGGGAGACCA | trRosetta | 180 | 130 |
| Q08 | Rous sarcoma virus | PKB174 | 128 | .........(((((((((((((............[[[[[[[[((((((..(((((......))))))))))))((((((....))))))......))))))))).))))))...........]]]]]]]]. | AAAUUUAUAGGGAGGGCCACUGUUCUCACUGUUGCGCUACAUCUGGCUAUUCCGCUCAAAUGGAAGCCAGACCACACGCCUGUGUGGAUUGACCAGUGGCCCCUCCCUGAAGGUAAACUUGUAGCGCU | trRosetta | 175 | 130 |
| Q09 | Methylosinus-1 | RF03108 | 175 | .(((.(((...((((((.....((((((....)))).))[[[[[[[[))))))))).)))....(((.(((((((((((.......))))((((((((....)))))))..))))))))))........(((((]]]]]]]].........(((....))).......)))))... | AGGGUGCGGUUUUCGAGAGAAGCGGGGCGGCUGCCCUCGUCCGCGCGCUCGAACGCUCCUCCGCCGGUCCGGCUUCUCCUUCGUGCGGAGGCCUUUGGAAACAAGGCGAAAGUCGGCCGAAUGUCCGAGGGUCGCGCGGGGAUAGACUACCCGAGAGGGGAAAACAACCCUGGC | FARFAR2 | 154 | 130 |
| Q10 | SCARNA2 | URS00008E39F0_9606 | 206 | .(((....(((((((((((.(((((((((((((((((..((((((((([[[[[[.............))))(((( | ACACAGGUGUGGGCUUACCUGACCAUGGCUGUCAAUAAGGAGGGGGGAGGUGUACUGGUAAUGAUCCCCAC | FARFAR2 | 161 | 130 |

| | | | | | | | | |
|---|---|---|---|---|---|---|---|---|
| | | | | ((((....)))))))).....((((<br>..............)))).(((((.((<br>(..........)))..)))))..)))<br>).....)))))))......)))))))<br>))..)))]]]]]].)))))))).. <br>.))). | AGCUUGUUGGUAAGCUGUUACAG<br>GAGCUUUAGGCAACACUUGCUCUG<br>GCCAUGCCCUUCAGUCCUGGCCCU<br>GGCUAUCUCUGUUCUGUUGGCGCA<br>UGAUGCCAUGGUUUUGGUCACCUC<br>AAGGCCUGCUUAGUGC | | | |
| Q11 | SV_g | spvincent | 240 | ..(((((((((([[[[[[[[.(<br>((((((((((........((((((<br>(((((((...........)))))))<br>)))))........)))))))))))<br>........))))))))))).(((((<br>(((((]]]]]]]](((((((((<br>.((............(((((((((..<br>..........))))))))))).......<br>.....)).))))))))).........<br>.))))))))))) | GACACAUCUCACACUCCUCCUGCC<br>UCACUCUGUUUAAAGGAGACGAG<br>GGUGCAAACCUCAAUUCUUAGUU<br>UGUACUUUCGCAGAAGAAAAACG<br>GAGUGAGAAUUUAUAUGUGAGAU<br>GUGACUAGCUUCCUUAGGAGGGG<br>UAUGCUCACAUUUUUCUUCUGCGG<br>AAGAUGUUUCAUAAGAAUUGAGG<br>AAACAUCUUAUACUUUUAUAAAG<br>AGUGAGUAUAGAGGAAGAGAAAG<br>GAAGCUAG | trRosetta | 161 | 130 |
| Q12 | SV_i | spvincent | 240 | ...((((((((((............[[<br>[[[[[[)))))))))....((((<br>((((((]]]]]]].[[[[[[[<br>[[[[)))))))))....((((((<br>((((]]]]]]]]]]]][[[[[[<br>[[))))))))))....((((((((<br>((]]]]]]]]..[[[[[[[[)<br>)))))))))....((((((((((]<br>]]]]]]]............))))))<br>))))....((((....)))).. | GGAAGUCUGUAGAAUUACUGUUA<br>AAUUCCCUUAUCUGCAGACUAUGA<br>AGUAGAGAUCUAAGGGAAUAAGG<br>UAGGAAUUGGUCUCUACUAUGAG<br>GGUUCUCUGAAUUCCUACCUUAAA<br>CGCGUACAGAGAACCCAGUAACAU<br>ACGUCGUACGCGUUUAAAAAUCUC<br>AUCGACGUAUGUAGUAAGUUAGU<br>GUGAUGAGAUUUUAAGGAGAGGA<br>ACACACUAGCUAAAUGGUUGAGA<br>AACCUA | trRosetta | 170 | 130 |
| Q13 | SV_h3 | spvincent | 204 | .((((((.......(......)......<br>...((((((((((((.[[[[[[[<br>[[[[[[[[............[[[[<br>[[............((((((((((((<br>(((((...(((((...........))<br>)))......)))))))))))))))<br>))))))))))))))..]]].]]]<br>..]]]]]]]]]]]]]]].))))))<br>). | AAUCACUAAGGUUUAGGGAAAUG<br>AAGAUGGGUGACUCAAUUCUUAG<br>AAUAUCGUAACGAUGAAGGAGUA<br>UAAUGUGAGGAUUUUCAUGUAAG<br>AUGUCACUAGCUUCCUUCUUACAU<br>GCAAAGUUCUCACAUGUUCCUUCA<br>AGGAAGCUAGUGACAUAAGAAUU<br>GAGUCAAUUCCAUCAACUAUCGUU<br>ACGAUAUUCAAGUGAUG | FARFAR2 | 133 | 130 |
| Q14 | SV_c | spvincent | 240 | .(((((((((((((((.......)<br>)))))))(((((((((((((...<br>....))))))))(((((((....(<br>((((((((((....((......((( | GUAGUCUGUGUCGUACACCGAGCC<br>UGUACGACCCUACUCGUAGUACUG<br>UAUGUGUGCUACGCGUGAGGUAA<br>GGAUGUGACUGAUUCUUACGCAU | FARFAR2 | 187 | 130 |

| | | | | | | | | |
|---|---|---|---|---|---|---|---|---|
| | | | | (((([[[[[[[[[[)))))))..)<br>)....)))))))))))....))....<br>..(((((((]]]]]]]]]])))))<br>))))))))((((((((.......))<br>))))))))))))((((((((....<br>...)))))))))))))))). | GCGUAGGUUUCCAGUCUAAAUCU<br>ACAAGUUUAUAUCAGUCACAUAU<br>GAACGCAUGCCAAUAGGUAGACU<br>GGACUUAUUGCUCACGGGAGCUCA<br>GGCUCGGUGAGCUUCAGUAGGACC<br>GGUACACAUACAGUACCGGUACAG<br>ACUAG | | | |
| Q15 | SV_f | spvincent | 240 | .((((((((((((((((((((...<br>......))))))))))))).(((((((<br>..((((((((.........))))))<br>)))((((((((((((((((((((<br>(.........)))))))))))).....<br>((([[[[[[[[[[[[[[[[...))<br>)...))))))))))(((((((((.<br>........)))))))))..))))))<br>)..((((((...]]]]]]]]]]]]]<br>]]])))))).)))))))))). | GAGAGUCUGUGUUGUACACUGAU<br>UGCUAACCAGUGUACAACAUAGU<br>ACUAGAUGUGACAGCUACGACACC<br>UGUCACGUGAGACUGCUCGUUAU<br>GCAUUGGUUAGCAAACAAUGCGU<br>AACAACAUGUUGCAAGUCCAACUU<br>CCAUGCGACAGAGAGUAGUCUCCG<br>GAGUUGCGUGUUGUAAGCAGUUU<br>CGAUAGUACUAUGCUAGUACCUG<br>GAAGUUGGACUUGUACUAGCACA<br>GACUCUA | trRosetta | 160 | 130 |
| Q16 | AK_PK240-3 | AndrewKa<br>e | 240 | .((((.(((((.(((((.....)))<br>)))))))......(((((((((..<br>...)))).))))))(((((((.((<br>(.....))))))))))...))))...<br>..(((((((((((((.....))))<br>)..((((((.....))))))(((.(<br>(((((.[[[[[[[[.))))))))<br>)..)))))))))...(((((((((<br>((..(((((.]]]]]]]].)))))<br>.......)))))))))))). | AGGACAGACUGACGGUCGAUUAG<br>ACUGCAGUCAAUUAAGAUCAGCU<br>ACGAUUAGUAGACUGGUCGAUCU<br>UCAGACGAUUAGUCGAGGAUCAU<br>AGUCCAAUAAGAUUAUAUGCUGG<br>CGAUUAGUCAGAAGAUGAGGAUU<br>ACUCGUCGAGACAUAUCAGGAUA<br>GGGAGAUAUGCUCAACAUGUAAU<br>CAAAGAUAAUGUUCCAAGAUUCA<br>CCCUAUCCAGAAUCAAUAUAAGGA<br>AUAUUAUCA | trRosetta | 164 | 130 |
| Q17 | pknot240 17 | mjt | 240 | ....(((((((((....[[[[...))<br>))))))).(((((((((.......)<br>))))))))..((((((.....))))<br>))...((((((.....[[[[..)))<br>)))..(((((((........))))))).<br>..(((((........)))))......<br>...((((......))))...(((((.<br>.......)))))..(((((.]]]]..<br>.))))).((((((((....]]]].))<br>))))).... | AAGAGUAUAUAGCACCAGGCCAA<br>GGCUAUAUACACACAUGUGUACG<br>ACAAACACAUGUGACGUAUGCAG<br>UAGGCAUACCGGGGCUCGAUGAG<br>AGGGCGCGAGCCAGGCCACGAAGC<br>CAGACGUGGCAGAGUGCCAUAAU<br>ACCGGCACAGGAAACCAAGCCCAU<br>AUAGGGGCCAUGGUCCAAGAAUA<br>UGGACCCAGCUACACCCUAGAGUA<br>GCAGGACACAAUGUGGCCAUGUG<br>UCCGAUA | FARFAR2 | 168 | 130 |

| | | | | | | | | |
|---|---|---|---|---|---|---|---|---|
| Q18 | Astros-Eli-mod | Astromon | 240 | ((((((((...((((((((((.[[[[[[[..)).)..)).))))))........(.(((((.((((((((((.[[[[.))))))))))...))))).)....))))))).((((((.(((((((((((..(((..(((]]]]...)).)))((((.(((((.((((.((((((....))))))).((((...(...]]]]]]])))))..))))))))))))))))))))))))))))))). | GGCAAAGGAUGAAGAGGACCGGAACUGACCAGCCAGCUGUCCCUCUUACCUAAAGACUUAAACCAAUGCCCUAGUGAGGGGGCAUUGGGCAUUAAGCCGUGACCUUUGCUAGAGAAUUAAAUGAGAUAAAAGAGGCCUCACGCAGGAUCUGGCAUAGAGGAGGUGAUCAGCAAAUGUUUGUUGAAAAGGUUUGACAGGUCAGUCCCUUCCCACCCCUCUUGCUUGUCUUAUUUAUUCUCA | FARFAR2 | 148 | 130 |
| Q19 | SV_r7_240_4 | spvincent | 240 | ((((((((((((.((((((((([[[[[..))))))))).(((((((((.(((((((((((((((((.((((((((((((......))))))))))).((((((((((((..]]]]].))))))))))).))))))))).((((((((((..[[[[[[.....))))))))))...))))))).(((((((((((.]]]]]].....)))))))))).))))))))).))))))))))) | UCUGUGGUCUGUAGGUAGAGCGAAGGAUAACGUUCUACCACCUCGACUCAGGUUAGAGGCAGCGGGAGGUUUGGGGGAAACAGCAUUUCCUCAAGCCAGGUCUUGUGUUUUAUCCUUAACACGAGGCCACCUGCUGCCACGUGGCUCGAAAUCCGGUGACAAUCGAGUUACGACGUCUGAUCACACAUCGAGCAAACCGGAAAAGAUGUUCGAUGUGAGAGUUGAGGAACAGACUACAGA | trRosetta | 150 | 140 |
| Q20 | Crazy Legs study 7 | tone | 240 | .......((......(((((..((..[[[[[[[........))..(...((((((...(((((((.(.((......))).....))).))))....))))))...).....]]]]]]])))))).......[[[[[[...((((((.(.....(..[[[[[[.)........[).))))))....{{{{{{.{....]........]]]]]]..)).....}.}}}}}}...]]]]]]... | AAGCGGUUCUUGAAAGCGCGAUGGGAUUGCUAAACAUGUUGCCAUGAGACUAGUCGAUGCGCAGACAUUGUUAAGUCAUGAAAAUCUAGCGCGAAUGAUUAGGAUUAAACGUUAGCGACGCGCAAAACAGUUUUCGUAAUGUACGGCGAAUCGAUUAUACGUCGAAUAUCGAGCUGUGCAUAAAGUUUGGACAGUAUCAAAACGAUGUAUAAAGAACUAAGUUCGAGCUGACGAAAAGUC | FARFAR2 | 119 | 140 |

*The target structure for the round 1 submission changed after the P06 Target opened. Both structures were used for computing OpenKnot score (maximum score assigned to designs).

**Table S5.** Targets for OpenKnot AI Rounds 3 and 4.

| | gRNAde (Mol9) PDB ID 10ZT EMD-75574 | MPNN-fixbb (Mol14) PDB ID 10ZU EMD-75575 | Struct2Seq-SHAPE (Mol13) PDB ID 11EH EMD-75648 | MPNN-RFdiff (Mol11) PDB ID 11AG EMD-75584 |
|---|---|---|---|---|
| **Data collection and Processing** | | | | |
| Microscope | Titan Krios G4 | Titan Krios G4 | Titan Krios G4 | Titan Krios G4 |
| Voltage (keV) | 300 | 300 | 300 | 300 |
| Camera | Falcon 4i | Falcon 4i | Falcon 4i | Falcon 4i |
| Magnification | 165,000x | 165,000x | 165,000x | 165,000x |
| Pixel size at detector (Å/pixel) | 0.7336 | 0.7336 | 0.7336 | 0.7336 |
| Total electron exposure ($e^{-}$/Å$^2$) | 50 | 50 | 50 | 50 |
| Exposure rate ($e^{-}$/pixel/sec) | 14.2 | 14.3 | 15.7 | 14.3 |
| Number of frames collected during exposure | 1134 | 1116 | 1116 | 1017 |
| Defocus range (µm) | -0.6 to -2.0 | -0.6 to -2.0 | -0.8 to -2.0 | -0.6 to -2.0 |
| Automation software | SerialEM | SerialEM | SerialEM | SerialEM |
| Energy filter slit width (if used) | 6 eV | 6 eV | 6 eV | 6 eV |
| Micrographs collected (no.) | 10,689 | 12,594 | 15,834 | 17,275 |
| Micrographs used (no.) | 10,689 | 8077 | 15,834 | 14,383 |
| Total extracted particles (no.) | 2,141,497 | 1,223,535 | 2,269,760 | 4,434,362 |
| Refined particles (no.) | 509,523 | 592,489 | 1,137,693 | 1,124,578 |
| Final particles (no.) | 479,673 | 113,232 | 969,545 | 155,837 |
| Point-group | C1 | C1 | C1 | C2 |
| Resolution (global, Å) | | | | |
| FSC 0.5 (unmasked/masked) | 4.8 / 3.2 | 6.4 / 4.7 | 6.8 / 6.0 | 7.8 / 6.8 |
| FSC 0.143 (unmasked/masked) | 3.6 / 2.9 | 4.8 / 4.0 | 5.2 / 4.8 | 5.4 / 4.8 |
| Resolution range (local, Å) | 0 – 22.3 | 1.42 – 21.2 | 0 – 9.5 | 0 – 11.5 |
| Resolution range due to anisotropy (Å) | 2.83 - 4.18 | 3.71 – 4.84 | 4.1 – 6.11 | 4.62 – 6.97 |
| Map sharpening *B* factor (Å$^2$) | -34.0 | -101.11 | -318.7 | -249.2 |
| Map sharpening methods | Local | Local | Global | Global |
| **Model composition** | | | | |
| RNA/DNA (nt) | 96 | 96 | 96 | 202 |
| **Model Refinement** | | | | |
| Refinement package | PHENIX | PHENIX | PHENIX | PHENIX |
| - real or reciprocal space | Real space | Real space | Real space | Real space |
| - resolution cutoff (Å) | 2.9 | 4.0 | 4.8 | 4.8 |
| Model-Map scores | | | | |
| - CC | 0.85 | 0.76 | 0.76 | 0.68 |
| - Average FSC (up to FSC 0.143) | 0.691 | 0.499 | 0.529 | 0.466 |
| *B* factors (Å$^2$) | | | | |
| RNA/DNA | 74.14 | 122.76 | 55.94 | 136.51 |
| R.m.s. deviations from ideal values | | | | |
| Bond lengths (Å) | 0.006 | 0.004 | 0.006 | 0.003 |
| Bond angles (°) | 0.829 | 0.693 | 1.039 | 0.617 |
| **Validation** | | | | |
| MolProbity score | 2.59 | 2.74 | 2.97 | 2.7 |
| Clashscore | 8.06 | 11.86 | 21.04 | 10.7 |
| **Primary contributors** | N.S.,D.B.H. | D.B.H.,N.S. | A.M.,J.H. | J.H.,N.S. |

**Table S6.** Cryo-EM data collection, refinement, and validation statistics.

| Name | Sequence |
|---|---|
| **UGAv3-5000-Tail2RC** | CCATCTCATCCCTGCGTGTCTCCGACTGCACGTCATGCAGTTGTTGTTGTTGTTTCTTT |
| **UGAv3-5001-Tail2RC** | CCATCTCATCCCTGCGTGTCTCCGACTGCACGAGATGCAGTTGTTGTTGTTGTTTCTTT |
| **UGAv3-5002-Tail2RC** | CCATCTCATCCCTGCGTGTCTCCGACTGCACGATGCGCAGTTGTTGTTGTTGTTTCTTT |
| **UGAv3-5003-Tail2RC** | CCATCTCATCCCTGCGTGTCTCCGACTGCACGATGCATAGTTGTTGTTGTTGTTTCTTT |
| **UGAv3-5004-Tail2RC** | CCATCTCATCCCTGCGTGTCTCCGACTGCACGCGCTGCAGTTGTTGTTGTTGTTTCTTT |
| **UGAv3-5005-Tail2RC** | CCATCTCATCCCTGCGTGTCTCCGACTGCACGCTGAGCAGTTGTTGTTGTTGTTTCTTT |
| **UGAv3-5006-Tail2RC** | CCATCTCATCCCTGCGTGTCTCCGACTGCACGCACAGCAGTTGTTGTTGTTGTTTCTTT |
| **UGAv3-5007-Tail2RC** | CCATCTCATCCCTGCGTGTCTCCGACTGCACGCATATCAGTTGTTGTTGTTGTTTCTTT |
| **UGB_trR2** | CTGTGTGCCTTGGCAGTCTCAGCTCAGACGTGTGCTCTTCCGATCTGGGAACGACTCGAGTAGAGTCGAAAA |
| **Eterna Forward Primer** | TTCTAATACGACTCACTATAGGGAACGACTCGAGTAGAGTCGAAAA |
| **Tail2 - RC Primer** | GTTGTTGTTGTTGTTTCTTT |
| **Starting sequence W03 gBlock** | GGGAACGACTCGAGTAGAGTCGAAAACGCGGAAACAATGATGAATGGGTTTAAATTGGGCACTTGACTCATTTTGAGTTAGTAGTGCAACCGACCGTGCTAAAACGTCTGATCTTCGGATCAGACGGGATTACGCTCCCTTCGGGGAGCGTAATCCAAAAGAAACAACAACAACAAC |
| **Eterna W03 gBlock** | GGGAACGACTCGAGTAGAGTCGAAAACGCGGACACAATGATGACTGCGTTTAAATTGGGCACTTGACGCAGTTTGAGTGAGTAGTGCAACCGACCGTGAAAAAACGTCTGATCTTCGGATCAGACGCGCGTGGTAGGTCTTCGGACCTACCACGCGAAAAGAAACAACAACAACAAC |
| **MPNN-fixbb W03 gBlock** | GGGAACGACTCGAGTAGAGTCGAAAAGGGGGAACCGAAGATGCAACGGCTATATTGGGACGTCTGGCTGTTGTTGAGGTGCTGACGCATCCAACCCTCTCAAAACGTCTGATCTTCGGATCAGACGGTGCACCGGATCGTTCGCGATCCGGTGCACAAAAGAAACAACAACAACAAC |
| **CdiGMP-Class I gBlock** | ACTCGAGTAGAGTCGAAAAGATGGACAATACAGTGATGTATTGTCCATCAAACTTGACTTGTCACGCACAGGGCAAACCATTCGAAAGAGTGGGACGCAAAGCCTCCGGCCTAAACCAGAAGACATGGTAGGTAGCGGGGTTACCGATGAAAAGAAACAACAACAACAAC |

**Table S7.** Oligonucleotides and gene fragments (gBlock) used for SHAPE chemical mapping experiments. Sequence coloring: T7 promoter – gold; 3´ constant Tail2 or Tail2 reverse complement – green; 5´ constant sequence – purple; barcode for Ultima genome analyzer – red.

| Name | Sequence |
|---|---|
| **gRNAde (Mol9)** | GUGCCACGGGGAGCGAUAGCUCCGGGUGAAAACCCUGGGGGCCGUGGCUAGGCUGGAGGCCCCC<br>GGCAGGGAGUGAAGGACGGAACAAGUAUGGCGUUCGCGCCAUGCUUGAACCACCGUAUACCGAA<br>CGGUACGUACGGUGGUGAAACAAACAAAUAAACUAAAUUAUGUGUGCCCGGCAUGGGUGCAGUC<br>UAUAGGGUGAGAGUCCCGAACUGUGAAGGCAGAAGUAACAGUUAGCCUAACGCAAGGGUGUCCG<br>UGGCGACAUGGAAUCUGAAGGAAGCGGACGGCAAACCUUCGGUCUGAGGAACACGAACUUCAUA<br>UGAGGCUAGGUAUCAAUGGAUGAGUUUGCAUAACAAAACAAAGUCCUUUCUGCCAAAGUUGGU<br>ACAGAGUAAAUGAAGCAGAUUGAUGAAGGGAAAGACUGCAUUCUUACCCGGGGAGGUCUGAGC<br>UUUCGAGCUCAGAAGUCAGCAGAAGUCAUAGUAACUCCCUGCCGUACACGAAAGUGUACCCUCC<br>AGCCGGAUC |
| **MPNN-fixbb (Mol14)** | GGACGCCCGGCGCUGUAGCGCCGGCCGAAAGGCCACGCCCAGGCGUCCUGGCGGGCCAGGGCGGU<br>CGGGGAGUGAAGGACGGAACAAGUAUGGCGUUCGCGCCAUGCUUGAACCACCGUAUACCGAACG<br>GUACGUACGGUGGUGAAACAAACAAAUAAACUAAAUUAUGUGUGCCCGGCAUGGGUGCAGUCUA<br>UAGGGUGAGAGUCCCGAACUGUGAAGGCAGAAGUAACAGUUAGCCUAACGCAAGGGUGUCCGUG<br>GCGACAUGGAAUCUGAAGGAAGCGGACGGCAAACCUUCGGUCUGAGGAACACGAACUUCAUAUG<br>AGGCUAGGUAUCAAUGGAUGAGUUUGCAUAACAAAACAAAGUCCUUUCUGCCAAAGUUGGUACA<br>GAGUAAAUGAAGCAGAUUGAUGAAGGGAAAGACUGCAUUCUUACCCGGGGAGGUCUGAGCUUU<br>CGAGCUCAGAAGUCAGCAGAAGUCAUAGUAACUCCCCGACGGGCCCGAAAGGGCCCAGGCCCGC<br>CGGAUC |
| **Struct2Seq-SHAPE (Mol13)** | GCCCUCUCUACACCUAUCGGUGUACCCAUAAGGGUCGCCCCAGAGAGGGCCCACUCACCGGGGCC<br>ACAGGGAGUGAAGGACGGAACAAGUAUGGCGUUCGCGCCAUGCUUGAACCACCGUAUACCGAAC<br>GGUACGUACGGUGGUGAAACAAACAAAUAAACUAAAUUAUGUGUGCCCGGCAUGGGUGCAGUCU<br>AUAGGGUGAGAGUCCCGAACUGUGAAGGCAGAAGUAACAGUUAGCCUAACGCAAGGGUGUCCGU<br>GGCGACAUGGAAUCUGAAGGAAGCGGACGGCAAACCUUCGGUCUGAGGAACACGAACUUCAUAU<br>GAGGCUAGGUAUCAAUGGAUGAGUUUGCAUAACAAAACAAAGUCCUUUCUGCCAAAGUUGGUAC<br>AGAGUAAAUGAAGCAGAUUGAUGAAGGGAAAGACUGCAUUCUUACCCGGGGAGGUCUGAGCUU<br>UCGAGCUCAGAAGUCAGCAGAAGUCAUAGUAACUCCCUGUGCCGCCCCAAAGGGCGGAGUGAGU<br>GGGGAUC |
| **MPNN-RFdiff (Mol11)** | GCGCCGGAGGCCCUAAAGGGCCAAGCGAAAGCUUAGGGGGACCGGCGCAGGGAUAGUACCCCCC<br>ACCGGGAGUGAAGGACGGAACAAGUAUGGCGUUCGCGCCAUGCUUGAACCACCGUAUACCGAAC<br>GGUACGUACGGUGGUGAAACAAACAAAUAAACUAAAUUAUGUGUGCCCGGCAUGGGUGCAGUCU<br>AUAGGGUGAGAGUCCCGAACUGUGAAGGCAGAAGUAACAGUUAGCCUAACGCAAGGGUGUCCGU<br>GGCGACAUGGAAUCUGAAGGAAGCGGACGGCAAACCUUCGGUCUGAGGAACACGAACUUCAUAU<br>GAGGCUAGGUAUCAAUGGAUGAGUUUGCAUAACAAAACAAAGUCCUUUCUGCCAAAGUUGGUAC<br>AGAGUAAAUGAAGCAGAUUGAUGAAGGGAAAGACUGCAUUCUUACCCGGGGAGGUCUGAGCUU<br>UCGAGCUCAGAAGUCAGCAGAAGUCAUAGUAACUCCCGGUGCGGGCCUAUAGGCCCGAACUAUC<br>CCGGAUC |

**Table S8.** RNA sequences for P20 designs characterized by cryo-EM, colored by the design regions (green); prepended G to enable transcription by T7 RNA polymerase and group II intron scaffolds for cryo-EM (black); and 3´ sequence derived from BamHI restriction site (blue).

**Movie S1.**

Cryo-EM maps and coordinates for P20 Kissing Multiloops designs by Struct2SeQ-SHAPE, MPNN-fixbb, and gRNAde, compared to AlphaFold 3 prediction. Dashed lines mark location of scaffold.

**Movie S2.**

Cryo-EM maps and coordinates of dimer formed by P20 Kissing Multiloops design by MPNN-RFdiff.
